# Model-based data analysis of tissue growth in thin 3D printed scaffolds

**DOI:** 10.1101/2021.03.25.436898

**Authors:** Alexander P Browning, Oliver J Maclaren, Pascal R Buenzli, Matthew Lanaro, Mark C Allenby, Maria A Woodruff, Matthew J Simpson

## Abstract

Tissue growth in three-dimensional (3D) printed scaffolds enables exploration and control of cell behaviour in biologically realistic geometries. Cell proliferation and migration in these experiments have yet to be explicitly characterised, limiting the ability of experimentalists to determine the effects of various experimental conditions, such as scaffold geometry, on cell behaviour. We consider tissue growth by osteoblastic cells in melt electro-written scaffolds that comprise thin square pores with sizes that we deliberately vary. We collect highly detailed temporal measurements of the average cell density, tissue coverage, and tissue geometry. To quantify tissue growth in terms of the underlying cell proliferation and migration processes, we introduce and calibrate a mechanistic mathematical model based on the Porous-Fisher reaction-diffusion equation. Parameter estimates and uncertainty quantification through profile likelihood analysis reveal consistency in the rate of cell proliferation and steady-state cell density between pore sizes. This analysis also serves as an important model verification tool: while the use of reaction-diffusion models in biology is widespread, the appropriateness of these models to describe tissue growth in 3D scaffolds has yet to be explored. We find that the Porous-Fisher model is able to capture features relating to the cell density and tissue coverage, but is not able to capture geometric features relating to the circularity of the tissue interface. Our analysis identifies two distinct stages of tissue growth, suggests several areas for model refinement, and provides guidance for future experimental work that explores tissue growth in 3D printed scaffolds.

**Author Summary:** Advances in 3D printing technology have led to cell culture experiments that realistically capture natural biological environments. Despite the necessity of quantifying cell behaviour with parameters that can be compared between experiments, many existing mathematical models of tissue growth in these experiments neglect information relating to population size. We consider tissue growth by cells on 3D printed scaffolds that comprise square pores of various sizes in this work. We apply a relatively simple mathematical model based on the Porous-Fisher reaction-diffusion equation to interpret highly detailed measurements relating to both the cell density and the quantity of tissue deposited. We analyse the efficacy of such a model in capturing cell behaviour seen in the experiments and quantify cell behaviour in terms of parameters that carry a biologically meaningful interpretation. Our analysis identifies important areas for model refinement and provides guidance for future data-collection and experimentation that explores tissue growth in 3D printed scaffolds.

## 1 Introduction

Cell culture scaffolds provide biomimetic experimental models to explore tissue growth in essential biological processes such as bone remodelling and development [1–3]. Achieving control over tissue growth through these scaffolds has clinical applications such as replacing synthetic grafts with artificially regenerated tissues [1,4,5]. Three-dimensional (3D) printing technology [6–9] enables precise control of scaffold geometry, including the size and shape of the pores that comprise each scaffold. Despite these technological advances, the effects of scaffold geometry on scaffold-level properties of tissue growth, such as the time for tissue to close or *bridge* scaffold pores, and individual-level properties, such as cell proliferation and migration rates, are yet to be explicitly understood.

A preference away from traditional *in vitro* 2D culture and towards mimicking biological features, such as the bone micro-environment, through 3D scaffolds has been aided by more accurate 3D printing processes [6,7]. Technologies based on melt electrowriting [8,9] enable precise control of scaffold geometry, ensuring consistency and reproducibility. There is a significant body of research that guides the material and physical properties of scaffold construction, but a comparative scarcity on the influence of scaffold architecture on cell and tissue behaviour. In fact, several recent studies suggest that tissue growth in pore infilling experiments is strongly curvature controlled [10–13], which implies that pore shape and size play a significant role in tissue growth [11,13] since the average curvature of a pore is a function of its size [14].

In this work, we consider tissue growth by osteoblastic cells in a 3D printed scaffold formed of thin square pores with depth ≈100μm and side lengths ranging from 300 to 600μm (Fig. 1*a-d*). This thin geometry means that we can approximate the three-dimensional tissue growth as a depth-averaged two-dimensional phenomena [15]. Initially located only on the scaffold fibres, cells migrate and proliferate to form new tissue that bridges each pore over an experimental duration of 28 days (Fig. 1*e-h*). Scaffolds are systematically harvested and stained to obtain fluorescent microscopy images that provide highly detailed information about the pore bridging progress (which we measure as the proportion of the pore containing tissue) and the cell density within each pore. The variability in pore bridging we see in the experimental data is striking: smaller pores appear, on average, to bridge at earlier times (Fig. 1*e-f*) [14]; and some, but not all, larger pores are bridged at the conclusion of the experiment (Fig. 1*g-h*). While we expect larger pores—which require the development of a larger amount of tissue and cells to migrate a greater distance—to bridge more slowly [14], it is unclear whether there are also changes in cell behaviour between pore sizes. We aim to determine whether there are fundamental differences in cell proliferation and migration between different pore sizes while demonstrating the value of collecting experimental data relating to both tissue coverage and cell population.

**Figure 1.**
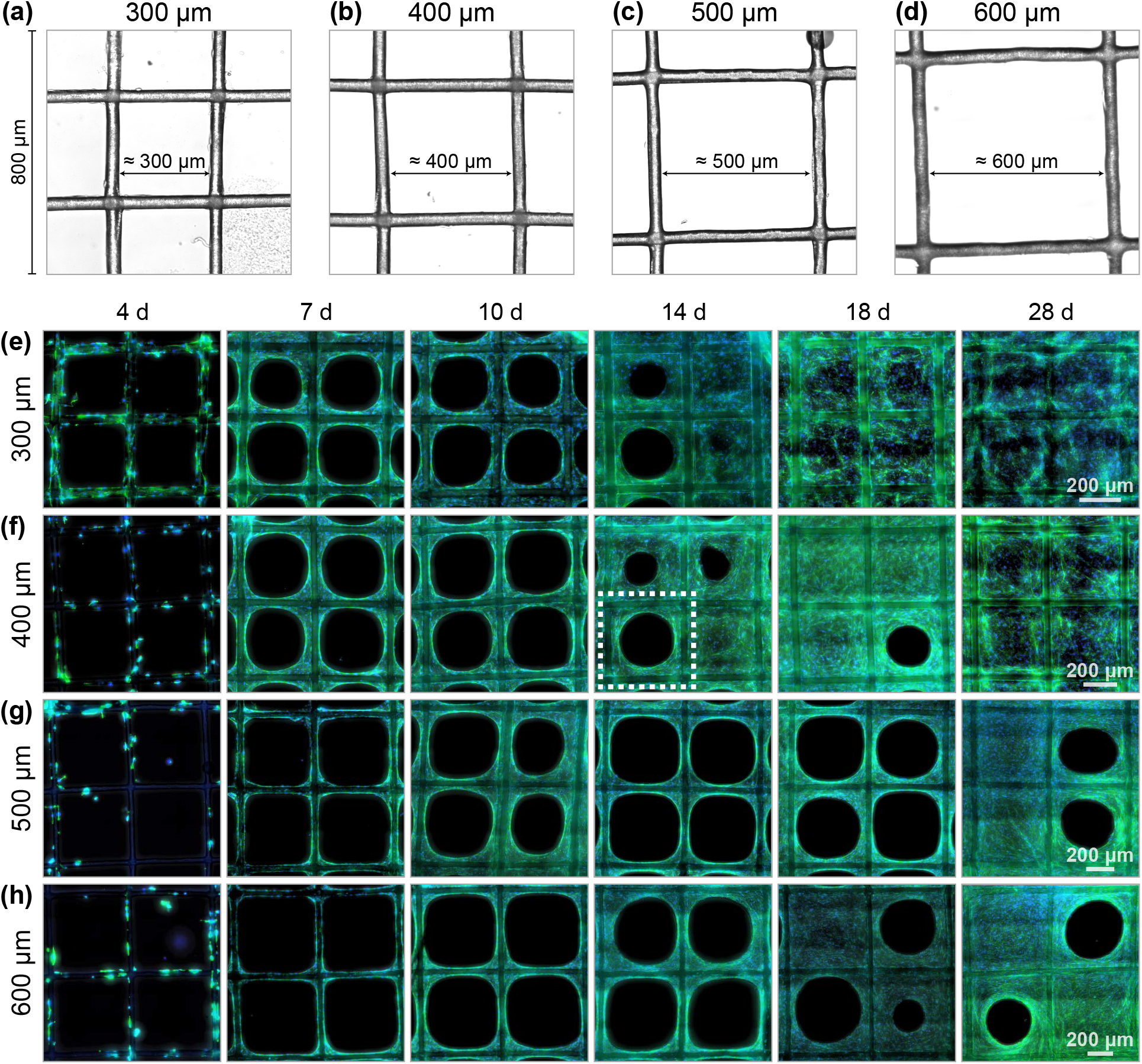
Scaffold geometry and experimental data. (a–d) Scaffolds comprise a grid of square pores with lengths ranging from 300 to 600 μm. Shown is a 800 × 800 μm DIC image taken from the central region of a scaffold for each pore size. (e–h) Composite fluorescence microscopy images of pore bridging experiments. Cell nuclei, stained with DAPI, are shown in the blue channel; tissue and cytoskeleton, stained with phalloidin, are indicated in the green channel. Scale varies between pore sizes, but is identical within a pore size and is indicated in day 28 images. It is important to note that scaffolds are fixed to obtain images: data from successive time-points are independent experiments.

To disentangle the effects of cell proliferation and migration on tissue growth, we perform model-based data analysis using a deterministic, continuum, process model [16]. Existing continuum models of tissue growth within porous scaffolds typically neglect information relating to properties such as cell count or density [17, 18]. Instead, the time-evolution of tissue interfaces are described using techniques ranging from continuum mechanics [3, 18, 19] to curvature flow [10,12,20-24]. While these models often provide good agreement with geometric features in experimental data, they yield parameter estimates that are purely phenomenological and lack a clear biophysical interpretation. We describe pore bridging using a relatively simple two-dimensional reaction-diffusion equation, often referred to as the Porous-Fisher model [14, 25]. This choice naturally accounts for density-dependent behaviour expected in these experiments: contact inhibition limits cell proliferation in high-density regions, and contact stimulates cell migration, leading to co-operative tissue growth that is limited in regions of low cell density.

We take a summary statistic and likelihood-based approach to parameter inference [26] to identify parameters that characterise cell behaviour both individually within each pore size, and across all pore sizes simultaneously. In comparison to our previous work [14], we consider a temporal dataset that includes information about both cell density and bridging progress. To quantify the uncertainty associated with parameter estimates—which may be non-identifiable from the available information in the experimental data—we perform profile likelihood analysis [27,28], which facilitates the computation of approximate confidence intervals [29–31]. We compare parameter estimates that quantify cell proliferation and migration rates across pore sizes to determine whether pore size, and by extension, curvature, influence cell behaviour. For example, if pore size and, by extension, curvature, play a significant role in cell proliferation, we would expect the estimates of the cell proliferation rate to vary significantly between pore sizes.

Compared to models of 2D culture, which are well developed and routinely applied in experimental design [16,32-36], there is little data-based modelling guidance for tissue growth within 3D scaffolds. Development and verification of mechanistic models for pore bridging is essential: models can guide engineering design choices in scaffold construction to optimise and control tissue growth [37]. Despite the widespread application of reaction-diffusion models in collective cell behaviour [32,38-40] and biology more broadly [41–44], their suitability to describe geometrically-induced phenomena—such as that arising from corners and the relatively small, constrained, domain in our experiments—remains largely unexplored. Qualitatively, the Porous-Fisher model produces results that capture key behaviours in the experimental data; namely both an increase in cell density over the duration of the experiment, and sharp-fronted tissue growth that bridges each pore (Fig. 2). A key focus of our work is to further verify the appropriateness of the Porous-Fisher model by comparing features not used for calibration to model predictions, and comparing parameter estimate and model behaviour across pore sizes. Given that tissue growth is thought to be curvature controlled [11], we focus on comparing geometric features in the data, such as circularity, to model predictions. Comparing parameter estimates and model predictions across pore sizes is crucial for model verification: if only a single experiment condition is considered, the model might appear to match the experimental data but be incapable of matching data across multiple experimental conditions without significantly varying the parameters [35,45]. Through this analysis, we identify several avenues for both future experimentation and model refinement.

**Figure 2.**
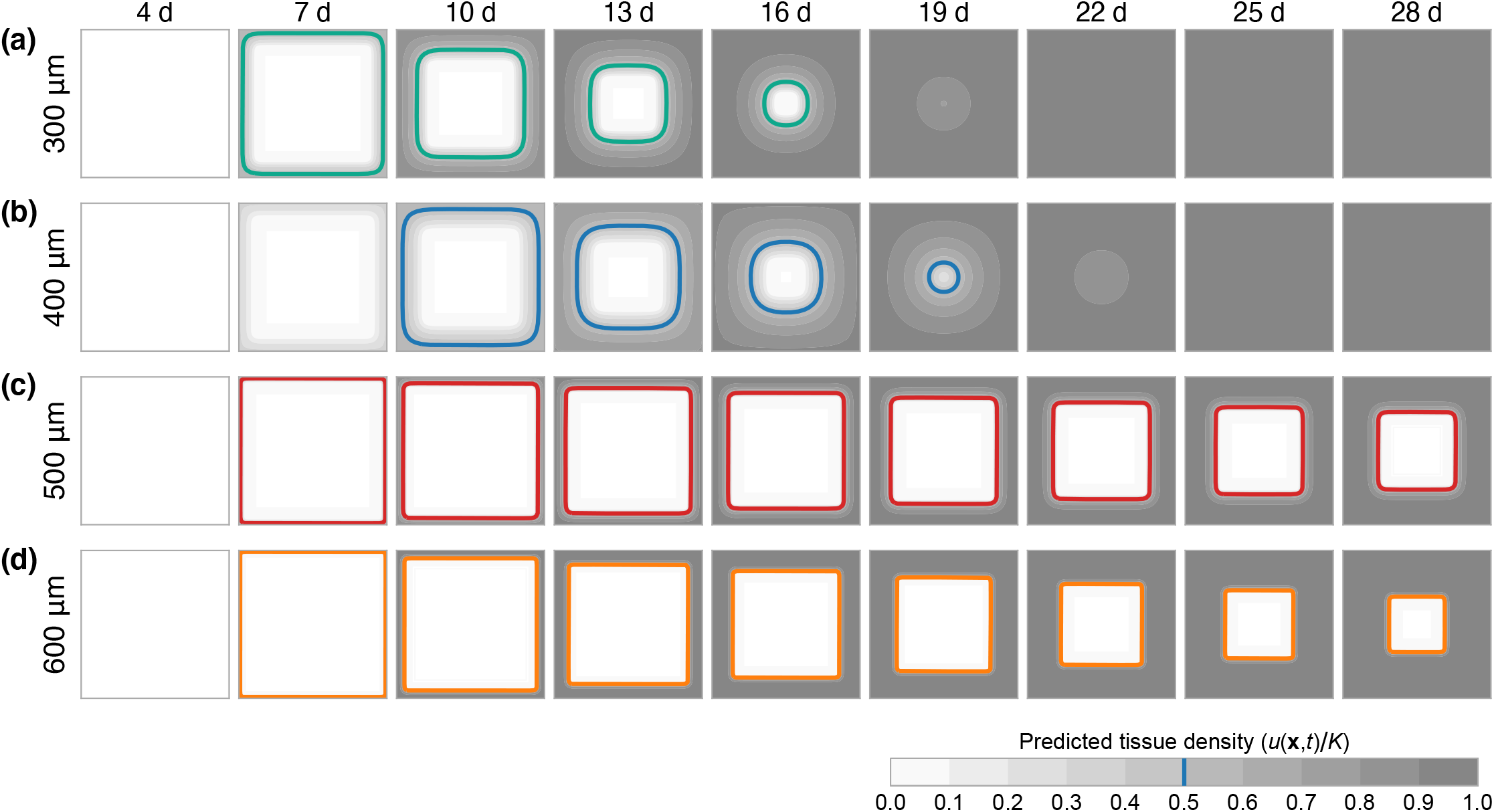
Model simulated tissue growth. Model simulation using the maximum likelihood estimate where information relating to cell density and tissue coverage are included in the likelihood. The coloured curves show the boundary of the ECM, taken to be τ = 0.5 (50%) of carrying capacity, *K*. Shown in greyscale is the density as a proportion of carrying capacity.

The outline of the work is as follows. We first describe the experimental model and methods used to summarise the data (Section 2.1). The data are available on GitHub as supplementary material. In Section 2.2, we describe a two-component mathematical model comprising both a deterministic process model that describes pore bridging dynamics and a probabilistic observation process that connects model predictions to noisy experimental observations. Subsequently, we outline the techniques used to obtain maximum likelihood estimates and likelihood profiles (Section 2.3). We present and discuss the results in Section 3 before outlining future experimental and mathematical modelling recommendations in Section 4. Code to reproduce all results are provided in the high-performance, open-source, Julia language on GitHub.

## 2 Methods

### 2.1 Pore bridging experiments

Polycaprolactone fibres of diameter 50 μm are fabricated into a two-layer scaffold of size 7 × 7 mm through melt electrospinning. The resultant scaffold has an overall thickness of approximately 100 μm (two fibre layers) and comprises square shaped pores of lengths 300, 400, 500 and 600 μm (Fig. 1*a-d*). Prior to cell seeding, scaffolds are sterilised and incubated in 5% CO_2_ overnight.

Murine calvarial osteoblastic cells (MC3T3-E1) [46] are cultured in α-MEM, 10% fetal bovine serum, and 1% penicillin-streptomycin (Thermo Fisher). Scaffolds are placed on top of non-adherent 2% agarose to prevent cell-to-plate attachment within a 48-well plate. Cells are detached using 0.05% trypsin and seeded at 7500cells in 250 μL media onto each scaffold within a 48-well plate (Nunc, Thermo Fisher). Cells are allowed 4h to attach to each scaffold before an additional 250 μL of media is added. Cell-seeded scaffolds are cultured in a humidified environment at 37 °C in 5% CO_2_ for 28 days. Media is changed every 2-3 days from day 1 to 14, every 1-2 days from day 15 to 21, then every day from day 22 to 28. Cell viability is assessed at day 10, 14 and 28 using calcein AM (to stain live cells) and ethidium homodimer (to stain dead cells).

**Figure 3.**
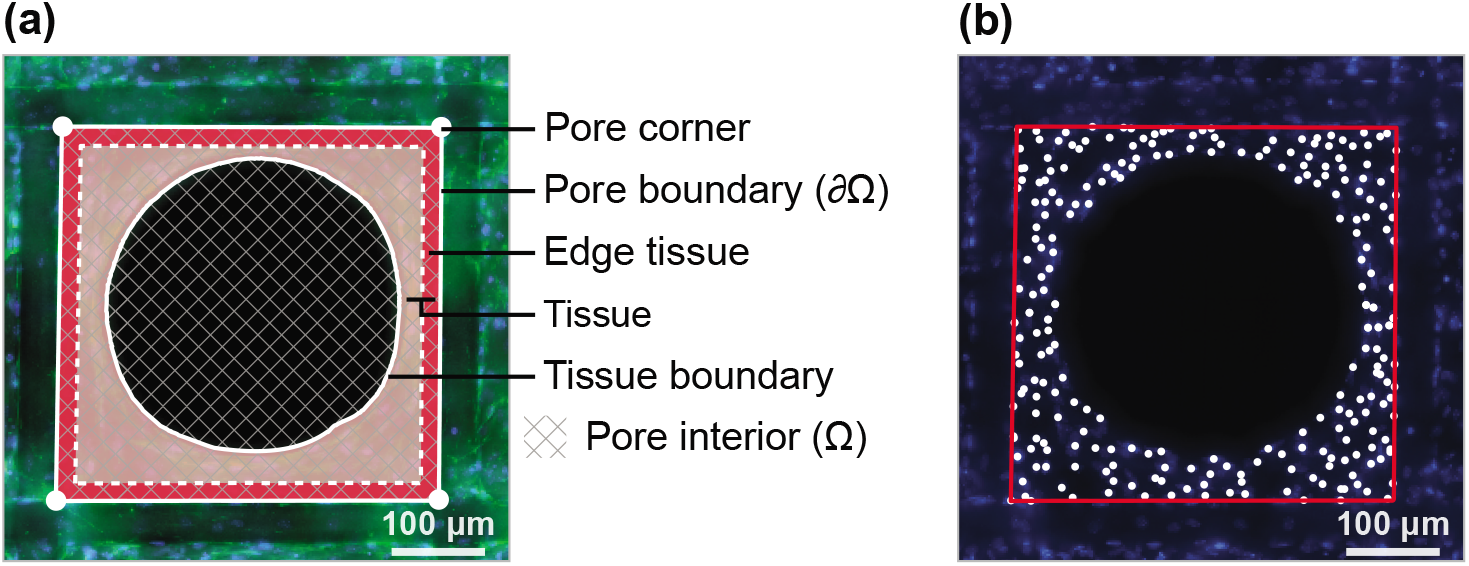
Data processing technique and experimental domain. Example data summarisation for a 400 μm pore at day 14, indicated in Fig. 1*f*. (a) The pore boundary and tissue identified using the semi-automated data processing approach. Also shown is the region classified as edge tissue. In the model, we denote the boundary of the pore ∂Ω, and the interior of the pore Ω. (b) DAPI image, showing cell nuclei, with the pore boundary and cell locations superimposed.

Cell-seeded scaffolds are fixed with 4% paraformaldehye at day 4, 7, 10, 14, 18, and 28. Replicates are stained with both DAPI and Alexa Fluor™ 488 Phalloidin (Thermo Fisher), which stain cell nuclei and actin filaments, respectively. Fluorescent microscopy (Leica AF6000 LX) is used to capture high resolution images of the centre of each scaffold. To accurately identify scaffold geometry, a differential interference contrast (DIC) image is also captured. Fixation, staining and microscopy are repeated across two or three replicates for each pore size and time point. Each experimental replicate yields information about 9 to 12 pores, providing tissue growth data across days 4 to 28 from 618 pores in total. In Fig. 1*e-h* we show composite images of four pores for each pore size, for each time point.

#### 2.1.1 Data summarisation

The tissue growth data are processed in a semi-automated fashion using MATLAB [47] to obtain information about the cell population and bridging progress in each pore (Fig. 3). First, the four corners of each pore are identified manually from the DIC image and thresholding is applied to the phalloidin image to establish the region in each pore containing tissue (Fig. 3*a*). Next, the location of the cells within each pore are identified using the thresholded DAPI image, which colours cell nuclei (Fig. 3*b*). Data are discarded for pores that are not deemed sufficiently regular in shape, or for which accurate measurements cannot be taken.

We summarise the experimental data obtained from each pore with four summary statistics, denoting 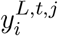 the *j*th observation of the *i*th summary statistic at time *t* for a pore of size length *L*. These are as follows.

1. Average cell density:

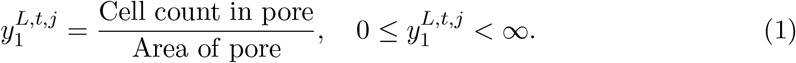
2. Coverage:

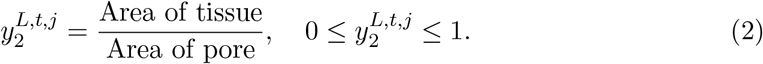
3. Edge density:

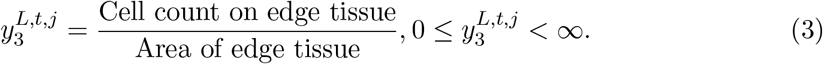 Here, we define edge tissue as tissue located within approximately 20 μm of the pore boundary (Fig. 3*a*).
4. Circularity:

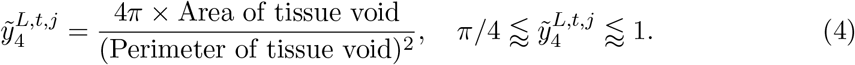

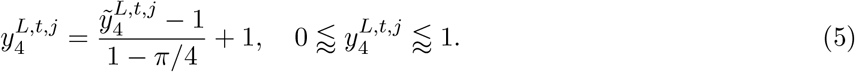 Here, 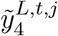 represents the standard measure of circularity or roundness [47], which approaches unity as the tissue void approaches a perfect circle. For a square shape, 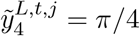. Since our experiments consider a scaffold that is approximately square, we normalise 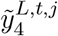 to obtain 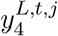 that still tends to unity as the tissue void approaches a perfect circle, but tends to zero as the tissue void approaches a square. To smooth out small-scale irregularities in the identified tissue shape, the convex hull of the largest contiguous tissue void is used to calculate the circularity [47].

### 2.2 Mathematical model

We interpret the pore bridging experiments with a deterministic spatio-temporal process model that aims to capture the key biological processes involved in tissue growth. To account for variability in the experimental data, we model experimental observations as normally distributed about predictions made through the process model [48–50]. In this section, we describe the process model and the probabilistic observation process used for analysis.

#### 2.2.1 Process model

The substrate of the wells containing the scaffolds is coated with non-adherent agarose that does not allow cell attachment, while cells are initially adhered to the scaffold fibres. Cells, therefore, cannot move freely into the pore void. Rather, cells work together to bridge the pore through interconnecting material such as extracellular matrix and intracellular actin filaments. Therefore, traditional models of cell migration based upon linear diffusion, which do not capture the sharp tissue boundary seen in the experimental data (Fig. 1*e-h*), are inappropriate.

In this work, we assume that cells move at a rate proportional to their own density and proliferate logistically to a maximum density of *K*, which we model with the Porous-Fisher [39] equation, given by

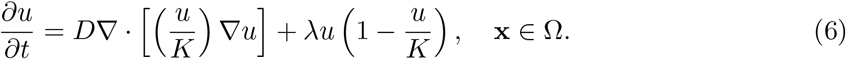

Given that the vertical depth of the pores is small compared to the horizontal length scale, and that we observe cells forming a thin horizontal layer of tissue that bridges each pore, we implicitly integrate out the vertical dimension [15] so that **x** = (*x,y*) and *u*(**x**, *t*) is a depthaveraged density, which we refer to as the cell density. In Eq. 6, Ω the interior of the pore (Fig. 3*a*) and ***θ*** = (*D, λ, K*) are parameters that relate to the diffusivity, proliferation rate, and carrying capacity, respectively.

The pore is surrounded by a fibre on which cells are initially placed approximately uniformly so that, on the fibre, ∇*u* = 0. We assume that both the proliferation rate and maximum packing density is the same as in the pore interior. Substituting ∇*u* = 0 into Eq. 6 recovers a time-dependent Dirichlet boundary condition on the edge of the pore

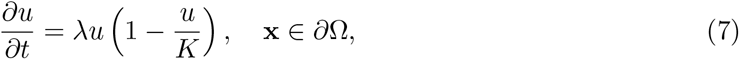

where ∂Ω represents the pore boundary (Fig. 3*a*).

Initially, cells appear distributed exclusively on the fibre, and not in the interior of the pore. It is not until after *t*_0_ = 4d that cells visibly start the pore bridging process (Fig. 1*a-d*). We, therefore, assume that at *t*_0_ = 4d, cells are distributed around the pore boundary (i.e., on the fibre) with an initial density *u*_0_, which we assume to be unknown and, therefore, estimate for each pore size. The initial condition is given by

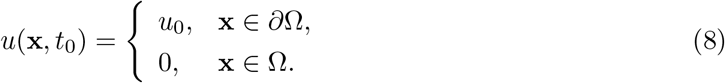

We solve Eq. 6-8 using a finite difference scheme based upon a discretisation with 101^2^ mesh points for each pore size. Due to the symmetry of the problem, we only solve Eq. 6-8 on a quarter-domain. To integrate the resultant system of ordinary differential equations, we apply the standard `Tsit5` routine in Julia [51,52]. Full details are available in the supplementary material.

#### 2.2.2 Observation process

Whereas output from the mathematical model is deterministic and comprises the cell density, *u*(**x**, *t*), as a function of space and time, the experimental observations comprise noisy observations of four summary statistics, 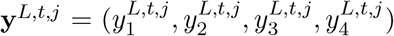. To compare model realisations to experimental observations, we define functions that map *u*(**x**, *t*) to summary statistics that correspond to those that summarise the experimental data. These functions are as follows.

1. Average cell density:

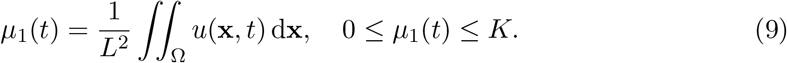 We approximate the integral in Eq. 9 numerically using the trapezoid rule.
2. Coverage:

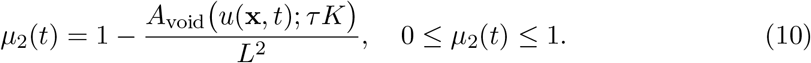 Here, *τ* represents a proportion of maximum cell density, *K*, at which tissue becomes visible, so that in regions where *u*(*t, x, y*) > τ*K*, cells are considered part of the observed newly formed tissue and A_void_ (*u*(**x**, *t*); τ*K*) is the area of the tissue void. In this work, we fix τ = 0.5, so that the tissue boundary in the model is assumed to be where the density is 50% of the maximum [14]. To calculate the area of the tissue void, A_void_(*u*(**x**, *t*); τ*K*), we apply an interpolation method to approximate the tissue boundary (supplementary material). This approach ensures that *μ*_2_(*t*) remains a continuous function in the parameter space, which is desirable for computational inference.
3. Edge density:

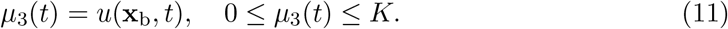 Here, **x**_b_ is any point on the pore boundary (the modelled cell density is homogeneous on the pore boundary); we set **x**_b_ = (0, 0).
4. Circularity:

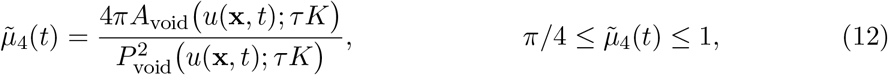

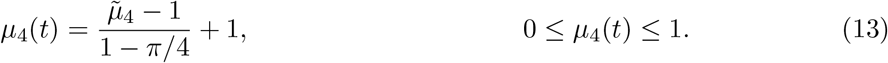 Here, *P*_void_(*u*(**x**, *t*); τ*K*) is an interpolated approximation of the perimeter of the tissue boundary (supplementary material). As for 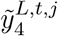, we normalise 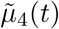 to obtain 0 ≤ *μ*_4_(*t*) ≤ 1 (Eq. 5). For simulations where the coverage exceeds 0.99, we set *μ*_4_(*t*) = 1 for convenience.

To account for biological noise and measurement error, we assume that model realisations describe the *expected behaviour* and that observations of the summary statistics are independent and normally distributed [48]. Therefore,

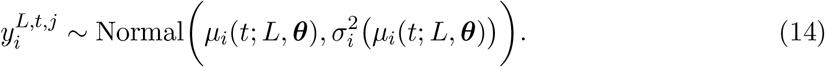

Here, we write *μ_i_*(*t*) = *μ_i_*(*t; L, **θ***) to emphasise the dependence of model realisations on the pore size, *L*, and set of unknown parameters, ***θ*** = (*D, λ, K*, *u*_0_). We observe in Fig. 4 that the variability in the experimental data varies significantly between both summary statistics and observation times. Therefore, we pre-estimate a variance function, *σ_i_*(*μ_i_*) as a function of the mean [48]. Here, we take *σ_i_*(·) to be a quadratic, with intercept of 10% of the maximum standard deviation observed for the summary statistic (supplementary material).

**Figure 4.**
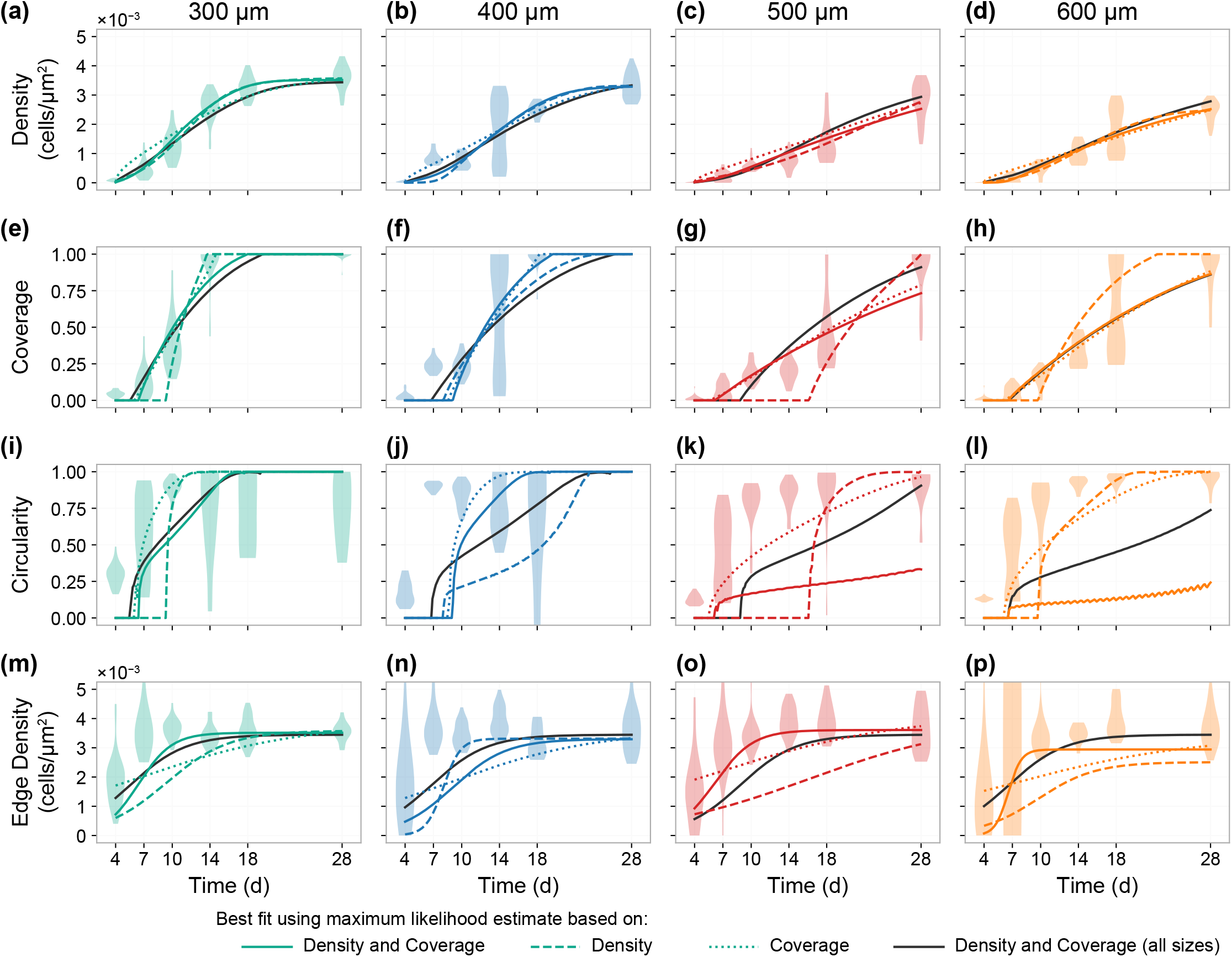
Processed experimental data and model fits. Experimental data and model fit showing (a-d) the density, (e-h) the coverage, (i-l) the circularity, and (m-p) the edge density. Violin plots show the experimental data. In each case, a model prediction is shown based on the maximum likelihood estimate that includes information relating to the cell density (dashed colour); cell density and tissue coverage (solid colour); tissue coverage with day 28 density measurements (dotted colour); and cell density and tissue coverage from all pore sizes (solid grey).

### 2.3 Inference

We take a summary statistic, likelihood-based, approach to inference and sensitivity analysis. Given a set of observations from pores of size *L*, **Y**^*L*^ = {***y**^L,t,j^*}_*j,t*_, the log-likelihood function is given by

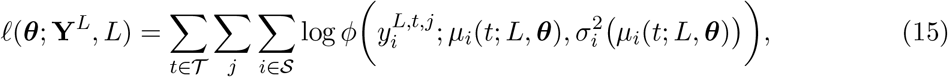

where 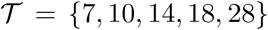 is the set of observation times (*t*_0_ = 4d is excluded from the analysis); 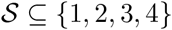 is the set of summary statistics included in the analysis; and *ϕ*(*x; μ, σ*^2^) is the normal density function.

#### 2.3.1 Parameter bounds

The set of unknown parameters, **θ** = (*D, λ, K, u*_0_), carry a physical interpretation so we can formulate realistic parameter bounds. The doubling time of MC3T3-E1 osteoblast cells in two-dimensional culture is approximately 15 h [46], which corresponds to a proliferation rate of approximately *λ* ≈ 1.1 d^-1^. Analysis based upon the overall bridging time of MC3T3-E1 osteoblast cells suggests D carries a magnitude of approximately 100 μm^2^ d^-1^ [14]. Results in Fig. 4a,b suggest that cell density is bounded above by approximately 4 × 10^-3^ cells μm^-1^, which corresponds to a packing density where a monolayer of cells occupy the same amount of space as a disk with diameter of approximately 18 μm. Based on these values, we choose conservative bounds such that

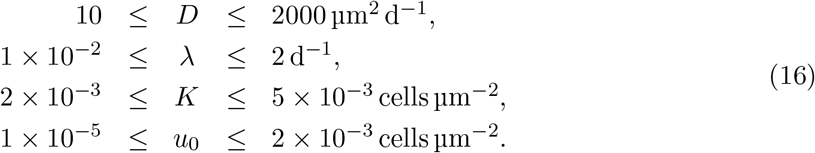

##### 2.3.2 Maximum likelihood estimation

We apply maximum likelihood estimation [53] to obtain a best fit parameter combination, 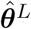, for each pore size. The maximum likelihood estimate (MLE) is given by

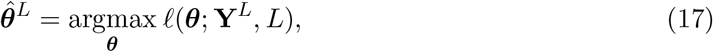

subject to the bounds given in Eq. 16.

To compute a numerical approximate the solution to Eq. 17, we employ both a global and local optimisation routine from the open-source `NLopt` optimisation library [54]. First, we run a global optimisation routine, based on the `DIRECT` algorithm [55], for a fixed amount of time (chosen to be 6 hours). This approach avoids the need to specify an initial guess of ***θ*** for the optimisation routine. We then use the output from the global optimisation routine as the initial guess in a the local optimisation algorithm `BOBYQA` [56]. We look for a maximum with absolute threshold of 10 ^−4^, several orders of magnitude below the threshold of 1.92 for an approximate univariate 95% confidence interval from a normalised-log-profile-likelihood [53].

##### 2.3.3 Profile likelihood analysis

While point estimates provide a means of assessing the ability of the model to capture features in the data, we are interested in establishing parameter uncertainties and comparing estimates across pore sizes. To do this, we profile the log-likelihood function for each parameter [29, 30].

First, we partition the parameter space into a parameter (or group of parameters) of interest, ψ, and nuisance parameters, **γ**, such that ***θ*** = (ψ, γ). The profile log-likelihood for the parameter ψ is given by

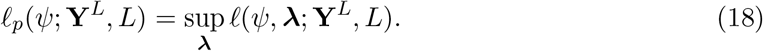

For example, to profile the diffusivity, we would specify ψ = *D* and γ = (*λ, K, u*_0_)· To obtain a value of *ℓ_p_*(*D; **Y**^L^, L*), we maximise the log-likelihood function in the case that *D* is fixed.

Likelihood-based confidence intervals can be defined from the profile log-likelihood by an asymptotic approximation using the chi-squared distribution, for sufficiently regular problems [53,57]. 95% confidence intervals and regions are given using the threshold values of 1.92 and 3.00 log-likelihood units below the maximum for univariate and bivariate profiles, respectively [53, 57, 58]. It is convenient to work with a normalised profile log-likelihood

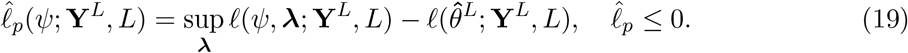

Here, a 95% confidence interval is given where 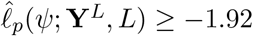, for example [53].

To compute numerical approximations to each profile log-likelihood, we employ the local optimisation routine `BOBYQA` [56]. The log-likelihood is profiled along a regular spaced grid, (ψ_1_,ψ_2_,…,ψ_M_), in series, starting at the grid point closest to the MLE, using the MLE as the initial guess [30]. Subsequent grid points use the output from the previous grid points as an initial guess. Again, we look for a maximum with absolute threshold of 10^-4^.

### 3 Results and Discussion

We interpret spatially-detailed, temporal, pore bridging data from a range of pore sizes using a relatively simple reaction-diffusion model. Our analysis considers data relating to the spatial characteristics of tissue growth—specifically, the tissue coverage and circularity of the tissue void—in addition to typical measurements, such as cell density. We aim to quantitatively determine whether there are fundamental differences in cell behaviour and tissue growth between different pore sizes, and verify the appropriateness of the reaction-diffusion model in explaining pore bridging, by comparing results across a series of experiments with various pore size. In particular, applications of reaction-diffusion models to describe tissue growth are typically limited to one-dimensional or unbounded geometries [32-36, 38-41]; there is comparatively little guidance on applying these models to describe the geometrically constrained phenomena we study.

In Fig. 1*e-h*, we show a subset of the experimental images obtained for each pore size over the duration of the experiment, and in Fig. 4 we show the summary statistics collected from the processed experimental data for each pore size. As each scaffold is fixed prior to staining and imaging, we note that data collected from successive time points are statistically independent. We work with average cell density (Fig. 4*a-d*) instead of cell count to allow direct comparison between pore sizes. It is not until after day four that cells migrate from the fibres into the pore void, so we exclude data collected at earlier time points from the analysis, and calibrate the model with observations taken after day four. Observations from day four itself are excluded as cells primarily occupy the fibres, which the model does not consider (Fig. 1*e-f*). By the end of the experiment (28 days), the majority of the 300 and 400 μm pores are bridged (85% and 100%, respectively), and the cell density appears very close to a steady-state (the average cell densities are 102% and 93% of the edge density, respectively). In comparison, several of the 500 and 600 μm pores do not bridge at the conclusion of the experiment (70% and 60% bridged at day 28, respectively), and, for these pore sizes, cell growth is more evident between days 18 and 28 (cell densities are 78% and 69% of edge density, respectively).

Using information about the cell density and tissue coverage, we calibrate the Porous-Fisher model to obtain a maximum likelihood estimate (MLE), ***θ**^L^*, individually for each pore size (Table 1). We show the solution of the model at the MLE, which we refer to as the best fit, along with the predicted tissue boundary in Fig. 2. Qualitatively, the behaviour predicted by the model matches that seen in Fig. 1 for the experimental data. First, the Porous-Fisher model predicts sharp-fronted migration, where regions ahead of the tissue boundary are devoid of cells. Second, we see cell migration drive tissue growth that bridges each pore. Pore bridging appears to occur at a slower rate for the larger pores, consistent with experimental observations. A counter-intuitive result that highlights the variability in pore bridging we see in the experimental data is that the 600 μm pores are predicted to bridge faster than the 500 μm pores: this is also seen in the experimental data, where at day 18 tissue coverage is greater in the 600 μm than the 500 μm pores (Fig. 1).

**Table 1.** Maximum likelihood estimates obtained by calibrating the Porous-Fisher equation to information relating to the cell density and tissue coverage. Asymptotic 95% confidence intervals, approximated using the profile likelihoods (Fig. 6) are given in parentheses. All values are stated to three significant figures.

| | $D$ ($\mu\text{m}^2 \text{d}^{-1}$) | | $\lambda$ ($\text{d}^{-1}$) | | $K$ (cells/$\mu\text{m}^2$) | |
| --- | --- | --- | --- | --- | --- | --- |
| 300 $\mu\text{m}$ | 397 | (290,653) | 0.561 | (0.353,0.858) | 0.00352 | (0.00338,0.00361) |
| 400 $\mu\text{m}$ | 1030 | (525,1690) | 0.35 | (0.191,0.694) | 0.0033 | (0.00314,0.00343) |
| 500 $\mu\text{m}$ | 117 | (40.6,269) | 0.497 | (0.238,1.21) | 0.00361 | (0.00322,0.00401) |
| 600 $\mu\text{m}$ | 99.9 | (54.7,240) | 1.41 | (0.621,2.0) | 0.00294 | (0.00271,0.00319) |
| All | 426 | (364,552) | 0.339 | (0.261,0.388) | 0.00345 | (0.00336,0.00355) |

In Fig. 4, we overlay a time-series of the best fit for each summary statistic with the experimental data, and in Fig. 5 we compare relationships between summary statistics predicted by the model to the experimental data. In all cases, we interpret realisations of the deterministic process model as the expected behaviour. To determine the distinct value of collecting information relating to the cell density and coverage, we also calculate the MLE in the case where we calibrate the model using (i) the cell density alone, and (ii) the coverage alongside day 28 observations of the cell density. Finally, to determine if the model can simultaneously match data across all pore sizes, we calculate the MLE using both cell density and tissue coverage information from all pore sizes (in this case, the initial density is allowed to vary between pore sizes). We show the best fits in these three additional scenarios in Fig. 4 and Fig. 5.

**Figure 5.**
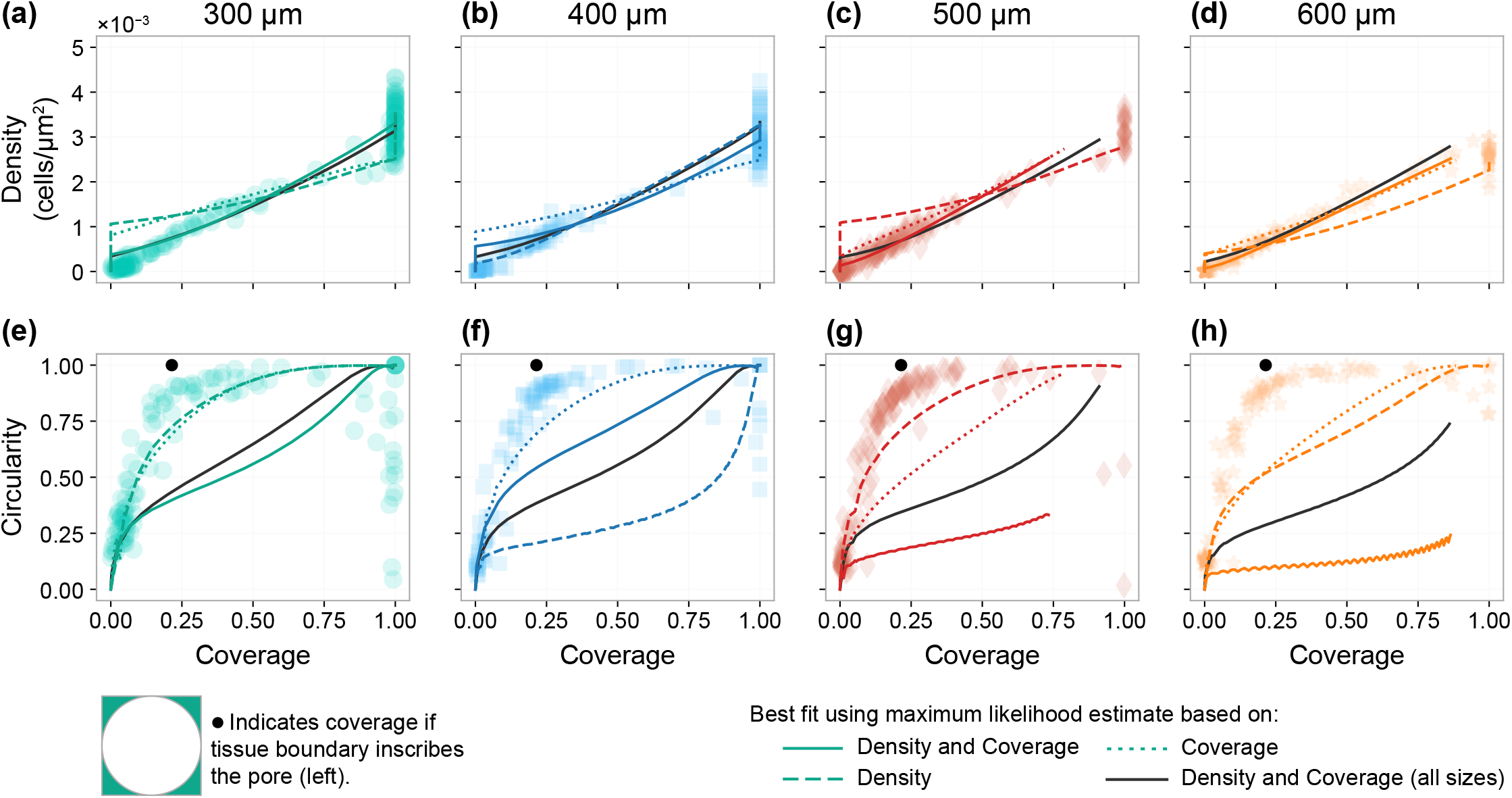
Relationships between experimental and simulated summary statistics. Experimental data and model fit showing the relationship between (a-d) tissue coverage and cell density, and (e-h) coverage and circularity. In each case, a model prediction is shown based on the maximum likelihood estimate based on the cell density (dashed colour); cell density and tissue coverage (solid colour); tissue coverage with day 28 density measurement (dotted colour); and cell density and tissue coverage from all pore sizes (solid grey).

Results in Fig. 4*a-h* show a clear value in considering information relating to tissue coverage. We see an excellent match with experimental observations of cell density for all pore sizes (Fig. 4*a-d*), even for cases where only tissue coverage and day 28 cell density observations are used for model calibration. Overall, we also see an agreement with experimental observations of tissue coverage; however, when the model is calibrated using cell density information alone, the best fit does not appear to capture early time tissue formation correctly (Fig. 4*e-h*). The model also provides an excellent match to experimental cell density and coverage observations when calibrated to all pore sizes simultaneously. These results are important as the model does not explicitly incorporate geometric behaviour (aside from the initial and boundary conditions) yet is still able to capture features relating to tissue coverage and cell density in the experimental data. This agreement between the model and experimental data is not only the case when parameter estimates are allowed to vary between pore sizes, but also when a single set of parameters is used to describe data across all pore sizes.

Comparison between model fits and experimental observations in Fig. 4*a-h* highlight how variable experimental observations are despite a large sample size of *n* = 618 pores: the average density and coverage for the 400 μm pores, for example, decreases by 12% from day 7 to 10 (the model monotonically increases), and observations at day 14 of the same pore size encompass observations at nearly every other time, (Fig. 4*b,f*). For this reason, we have excluded day 7 observations of 400 μm pores from results in the main text. In the supplementary material, we demonstrate that including these observations leads to results inconsistent with the other pore sizes. We address possible reasons for high levels of variability data later in the discussion.

A critical area in which the model fails to capture the experimental observations is in its ability to match the circularity of the tissue boundary in the larger 500 and 600 μm pores (Fig. 4*k,l* and Fig. 2*c,d*). We further verify this by calibrating the model to information that includes circularity, finding that the model best fit in this case does not match the circularity measurements seen in the experimental data (supplementary material). In Fig. 5*e-h* we explore the relationship between tissue coverage and circularity, which we note are both non-dimensional quantities and, therefore, can be directly compared between pore sizes. The relationships observed in the experimental data are remarkably consistent both between and within pore sizes, contrasting significantly to results in Fig. 4 that show highly variable observations. Comparing the tissue boundaries predicted by the model (Fig. 2) to the experimental data (Fig. 1) reveals why this may be the case. The model predicts initial tissue growth in both the corners and edge of the pore. In comparison, tissue growth in the experiments appears to occur initially only in the corners: it is not until the tissue boundary becomes almost completely circular, with a diameter equal to the pore size, that tissue growth occurs on the pore edge. We confirm this in Fig. 5*e-h* by calculating the coverage for a hypothetical, idealised, pore that forms a circular tissue void inscribed in the pore, equal to 100(1 — π/4)% for all pore sizes (indicated in black). We note that since the manufacturing process never leads to pores that are perfectly square, we do not expect to see a perfectly circular pore with coverage 100(1 — π/4)% in the experimental data. This corner corner bridging stage of tissue growth is not included in or captured by the Porous-Fisher model (Fig. 2). To develop a better understanding of corner bridging, we suggest future mathematical and experimental work focussed on corner bridging using scaffolds with pores large enough that tissues in adjacent corner tissues do not interact and start pore bridging [17, 22].

Point or maximum likelihood estimates for each parameter vary across pore sizes (Table 1), yet the model is able to match experimental observations of cell density and tissue coverage across all pore sizes with a single set of parameters (Fig. 4). To allow for parameter uncertainty when comparing parameters across pore sizes, we compute profile likelihoods (Fig. 6) and approximate confidence intervals (Table 1) for each parameter. Although profile likelihoods must be interpreted with care given that they depend not only on the process model but also the noise model, they provide valuable information about the sensitivity of the likelihood estimates we obtain. This is important as the point estimates provided by the maximum likelihood estimate give no information about parameter uncertainty, sensitivity or identifiability [29]. Although point estimates for each parameter appear to vary between pore sizes (Table 1), confidence intervals (Table 1) and likelihood profiles (Fig. 6) largely overlap, providing no evidence that these parameters vary across pore sizes. MLEs obtained for the diffusivity, *D*, from cell density information alone are much larger than we might expect, but examination of the profile likelihoods, which provide a lower, but no upper, bounded confidence interval, indicates that *D* is cannot be established unless information relating to tissue coverage is included. The largest discrepancy between pore sizes is seen in the diffusivity: estimates range from 400 to 1000 μm^2^ d^-1^, for the 300 and 400 μm pores, respectively (the larger variability and inconsistencies in data for the 400 μm pores leads to a much wider confidence interval than for the 300 μm pores), to below 200 μm^2^ d^-1^ for the 500 and 600 μm pores. This variability is consistent with estimates for cell diffusivities in two-dimensional culture, which often vary over several magnitudes across experimental conditions [59, 60].

**Figure 6.**
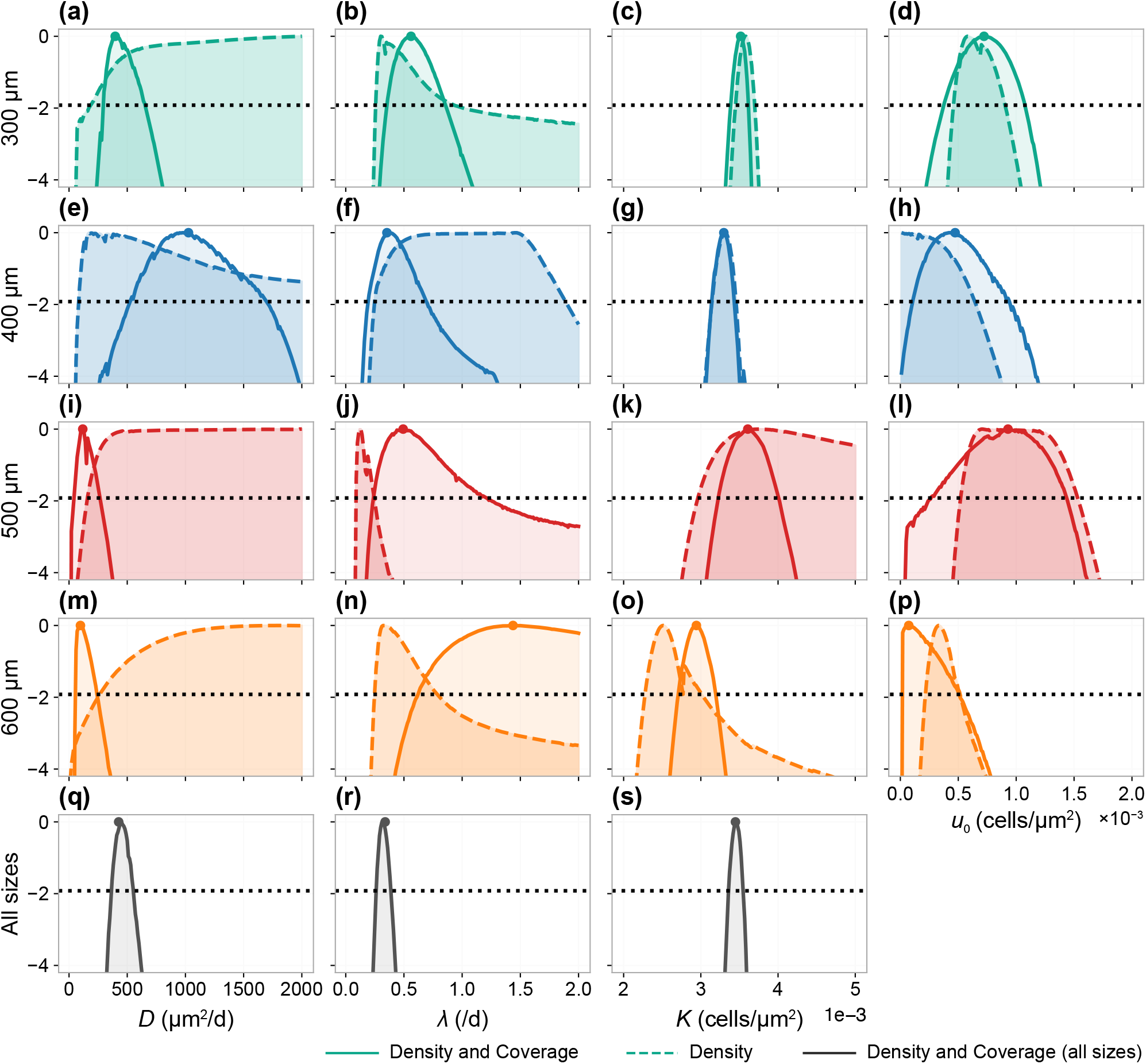
Profile likelihoods for calibrated model parameters. Profile likelihoods for each inferred parameter and pore size where only the density is used (dashed) and where both the density and coverage are used (solid). Dotted horizontal black line indicates the —1.92 contour which corresponds to an asymptotic 95% confidence interval for each parameter. Also shown are profiles for *D, λ* and *K* where information relating to cell density and tissue coverage information from all pores is included simultaneously.

Estimates, profiles and confidence intervals for the proliferation rate, λ, are remarkably consistent between pore sizes. While the model does not capture the shape of the tissue boundary, it does capture both the cell density and tissue coverage, suggesting that the crowding effects which lead to logistic growth in the experiments are also captured. In particular, our results in Fig. 6 suggest proliferation of MC3T3-E1 is similar between scaffolds of different sizes and is lower than a rate of λ ≈ 1.1 d^-1^ observed in two-dimensional culture [46] (this is also seen in Fig. 7). Another interesting result is the consistency in carrying capacity, K, of approximately 0.00345 (95% combined CI: (0.00336,0.00355)), which corresponds to an average packing density where a monolayer of cells occupy the same amount of space as a disk with diameter of 19 μm. An exception is for the largest 600 μm pore, which produces an estimate much lower than the other pores (95% CI: (0.0027,0.0032)). While this lower estimate may be consistent with average cell density observations (Fig. 4*d*), the higher estimate from the combined MLE is more consistent with behaviour at the edge of the pore (Fig. 4*p*). In some cases, the assumption of a constant carrying capacity across the entire pore may not be appropriate. It is not clear from the data alone whether this observation is due to actual variation in carrying capacity within a pore, or because net cell growth in the centre of the pore has not yet plateaued due to crowding effects. To answer this question, data must be collected over a longer experimental duration for these larger pores.

**Figure 7.**
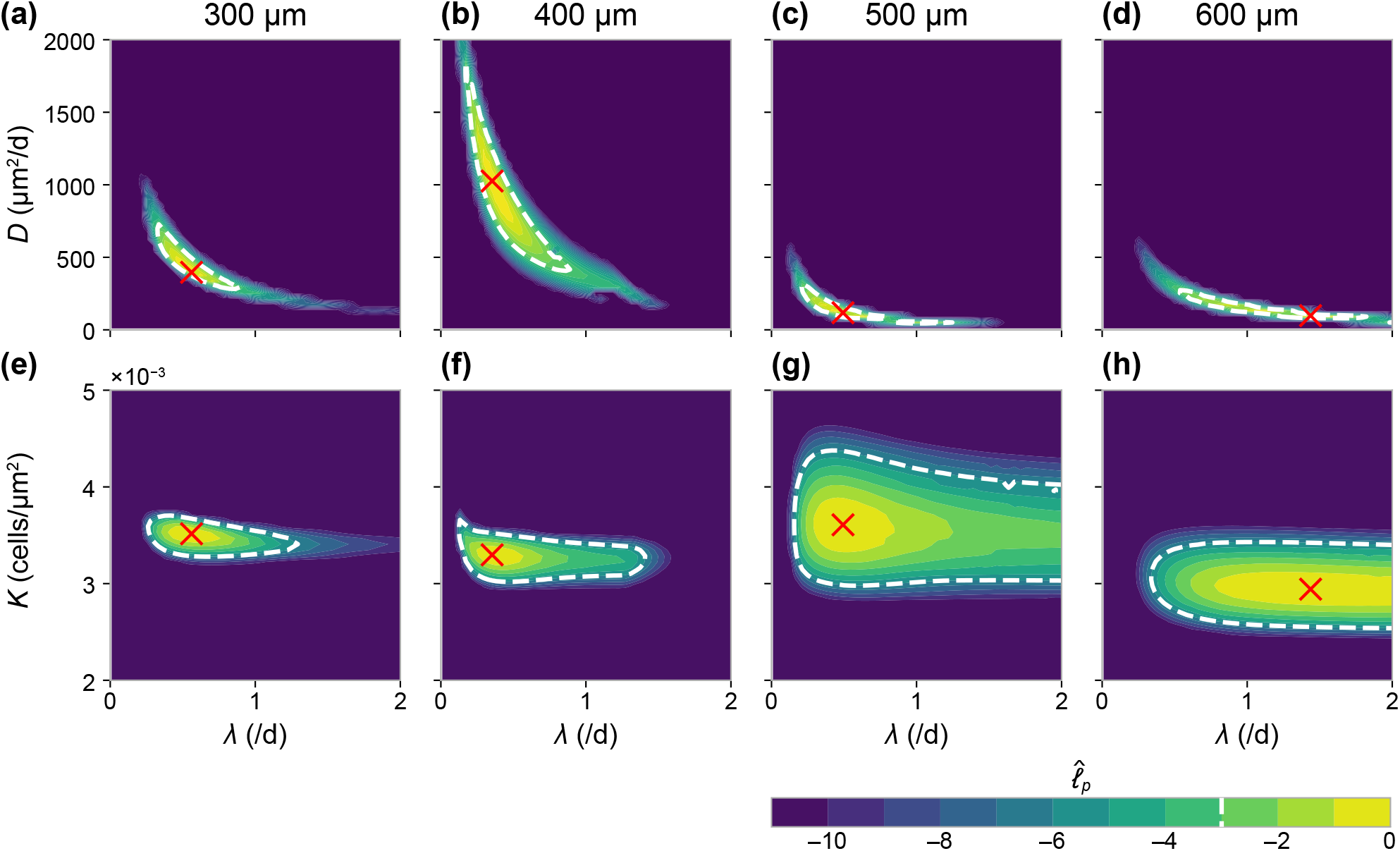
Bivariate profile likelihoods for calibrated model parameters. Bivariate profile likelihoods showing the relationship between estimates for (a-d) *D* and λ, and (e-h) λ and *K*. Dotted white lines indicates the —3.00 contours which corresponds to an approximate asymptotic 95% confidence region for each parameter combination.

In Fig. 7 we compute bivariate profiles to assess potential relationships between parameter estimates. First, examining the bivariate profiles between the proliferation rate, λ, and diffusivity, *D*, in Fig. 7*a-d*, reveals a hyperbolic relationship. This result is consistent with previous studies that establish only the product *D*λ using information about the position of the tissue interface [14,40], but that cannot establish individual values for these parameters. In our work, by using information relating to both cell density and tissue coverage, we are able to establish the individual values of *D* and λ within a region of compact support (a 95% confidence region is shown in Fig. 7*a-d*). Second, examining the bivariate profiles between the proliferation rate, λ, and carrying capacity, *K*, highlights the information obtainable from the 28 day experiment for each pore size. On average, the larger 500 μm and 600 μm pores do not bridge by the conclusion of the experiment, and we see comparatively large uncertainties in both the estimated proliferation rate and estimated carrying capacity (Fig. 7*g,h*). In contrast, results for the smaller 300 μm and 400 μm pores—the majority of which bridge by day 18—show that we are able to establish these parameters with a relatively small region (Fig. 7*e,f*). Although point estimates for the proliferation rate vary across pore sizes (Table 1), the bivariate profiles show a significant overlap in possible parameter values, indicating that these parameters are similar between pore sizes.

When the data are analysed as summary statistics that depend upon time, as in Fig. 4, we see a large amount of variability that cannot be fully captured by the observation noise process we define in Section 2.2.2. However, when relationships between summary statistics of each pore are analysed with respect to each other, independent of time, we see notably less variability (Fig. 5). These results suggest that time alone is a poor predictor of each summary statistic. In contrast, the dependence between summary statistics in Fig. 5 suggests that summary statistics have well defined relationships with relatively little variability. In the deterministic process model, the initial condition (which describes the expected value of each summary statistic on day 4) is taken to be a fixed density of cells distributed around the fibres. The majority of the variability in the temporal pore bridging data may be due to variability in the initial condition, which affects initial pore formation. For example, we expect tissue growth to be slower, or stationary, for pores that initially have a smaller density of cells distributed around the pore boundary at day 4. One way around the limitation of providing a homogeneous initial distribution of the cells in the experiments is to collect time-series data, where the same pore is imaged at multiple time points. There are two ways this information could be incorporated into the mathematical model. First, by including a time delay parameter for each data point that describes the delay until tissue formation inside the pore begins, that can be profiled out as a nuisance parameter in the analysis. Second, by capturing the variability directly by describing pore bridging as a differential equation where the initial density at the pore boundaries is a random variable.

Our results do not suggest significant differences in cell behaviour between pore sizes. Despite the Porous-Fisher model not explicitly incorporating geometric behaviour (aside from the initial and boundary conditions), we can capture information relating to both tissue coverage and cell density even when calibrated simultaneously to data from all pore sizes. By accounting for tissue coverage, we quantify a similar proliferation rate for all sizes based on a logistic growth assumption. The relationship between tissue boundary circularity and coverage is similar between all pore sizes. In all pores we see two stages of bridging: first, the corners bridge—this takes longer in the larger pores—and form an approximately circular tissue boundary; second, the pore closes and remains approximately circular in shape. These observations have also been made for triangular and hexagonally shaped pores [20], and convex pores [11]. Further experimental and modelling work is needed to disentangle the effect of each of these stages on overall pore bridging and tissue growth. For example, we suggest experimental work that investigates corner bridging and tissue establishment using non-constrained or “open” geometries [17,22], rather than the current geometry where tissue growth eventually closes a pore of finite size. To reduce overall variability in the data, variability in the initial condition should be accounted for through time-series imaging, where information about each pore is available at multiple time-points, and throughout each distinct stage.

Our thin three-dimensional experimental framework, and two-dimensional depth-averaged mathematical modelling framework, carry several advantages over more complex alternatives. In addition to information relating to tissue coverage, we are able to access detailed information about cell density, which we interpret with a mathematical model that quantifies cell behaviour with biophysical parameters such as proliferation and migration rates. This allows for comparison of cell behaviour between cell lines, allowing tissue growth optimisation with respect to cell line in addition to scaffold geometry. Our modelling framework is also extensible to co-culture systems that include multiple cell lines, which are more representative of *in vivo* tissue growth, through a coupled system of partial differential equations. Additionally, working with a thin three-dimensional experimental geometry reduces the need to account for additional extraneous factors on cell growth, such as nutrient availability. For example, typical *in vitro* three-dimensional tissue culture lack the vascular system that ensures homogeneous nutrient availability for *in vivo* tissue growth [3]. In comparison, our geometry results in a monolayer of cells that are all in direct contact with growth medium.

We suggest, in future, a hybrid modelling framework to describe each stage of pore bridging, rather than a single model that captures all stages of growth. While our analysis does not preclude generalisations of the Porous-Fisher model from capturing geometric features like circularity, reaction-diffusion models alone cannot account for both the corner bridging and pore closing stages of growth we see in the experimental data. Models based on continuum mechanics or curvature control have been successful in recapturing the initial stages of bridging seen experimental data [11,20,23], but typically neglect information relating to cell density. Once a circular tissue boundary is established, tissue growth may be quantified using density-dependent models such as those based on the Porous-Fisher equation, or agent based models [45, 61].

## 4 Conclusion and Outlook

We analyse experimental data from a series of pore bridging experiments using a relatively simple reaction-diffusion model based on the Porous-Fisher equation. In addition to commonly reported tissue coverage information, our model allows for the interpretation of information relating to cell density, and we see a clear value in considering both measurements. For example, the cell migration rate is often unidentifiable from information relating to cell density alone but becomes identifiable when information relating to tissue coverage is included. Compared to existing models of tissue growth that are largely phenomenological [17,18], our framework characterises cell behaviour with parameters that carry a biologically meaningful interpretation, such as cell proliferation and migration rates. We find no evidence to suggest that cell behaviour is dependent upon pore size. The cell proliferation rates, which are lower than that observed for two-dimensional culture, and carrying capacities are found to be remarkably similar across different pore sizes. This outcome suggests that our experimental protocols lead to consistent, reproducible tissue growth. This conclusion is not apparent without interpretation of the experimental data with a mechanistic mathematical model.

Our analysis identifies two distinct stages of pore bridging that are consistent between pore sizes: an initial corner bridging stage, and a latter hole closing stage. The Porous-Fisher model does not describe the initial corner bridging stage and, therefore, does not reproduce the shape of the tissue boundary. However, the model does match features relating to cell population and tissue coverage, thus capturing crowding effects and providing confidence in the estimated cell proliferation rates. We suggest that a better understanding of pore bridging can be formed through distinct theoretical models and experimental analysis that individually capture both the corner bridging and hole closing stages.

The experimental data used for model calibration suggests, at first, that pore bridging is a highly variable process. However, analysis of the relationships between summary statistics reveals this may not be the case. Rather, variability in both the initial distribution of cells on the scaffolds and corner bridging leads to a time-delay that cannot be accounted for with the information available from our data-collection method. These results highlight a potential value in designing an experiment to collect time-series observations, which will provide information about cell density and tissue coverage of each pore at multiple time points. This more detailed information will allow for the inclusion of more complicated mechanisms, such as directed migration through chemotaxis [43,62], mechanical effects at the tissue boundary [63,64], or the depletion of nutrients available to the cell population. At present, we find the complexity of the mathematical model is well suited to the level of information available in the experimental data, and we expect identifiability issues to arise if we were to interpret the current data with a more complex model.

Many of our conclusions could not have been made without considering data from multiple experimental geometries. The smaller pores, for example, give the impression that the model captures geometric features of pore closing; the inability of the model to capture these features is only evident when we analyse data for the larger pores. Comparing parameter estimates and profile likelihoods across experimental conditions is essential for constructing and verifying theoretical descriptions of pore bridging. Typical applications of mechanistic mathematical models to understand tissue formation usually involve working with a single experimental geometry, most often in a one-dimensional setting. These approaches cannot provide insight into the effect of high-dimensional geometric phenomena, such as corners, which we explore in our work.

In conclusion, our Porous-Fisher model successfully captures many of the key features of the experiments, providing a straightforward means of interpreting experimental observations in terms of the underlying cell proliferation and migration mechanisms that drive tissue growth. To the best of our knowledge, these mechanisms have never before been explicitly characterised for tissue growth in 3D-printed scaffolds.

## Data availability

Code and data used to produce the numerical results are available as a Julia module on GitHub at github.com/ap-browning/Pore-Bridging.

## Author Contributions

A.P.B. performed the data analysis, implemented the mathematical model, and wrote the paper. A.P.B. and M.L. processed the experimental data. M.L. and M.C.A. performed the experiments. All authors provided feedback and gave approval for final publication.

