## Supplementary Material Document for "Model-based data analysis of tissue growth in thin 3D printed scaffolds"

### Supplementary material for "Model-based data analysis of tissue growth in thin 3D printed scaffolds"

#### Contents

**S1 Numerical solution of the PDE** **2**

**S2 Interpolating tissue void boundary from PDE solution** **4**

**S3 Noise model** **5**

**S4 Including all $L = 400\,\mu\text{m}$ data** **6**

**S5 Inference using circularity** **7**

**S6 Fisher-Kolmogorov model** **9**

#### Code availability

This supporting material document is supplementary to, and refers to, code available on GitHub at github.com/ap-browning/Pore-Bridging.

#### S1 Numerical solution of the PDE

We write the partial differential equation (PDE) model as

$$
\begin{aligned}
\frac{\partial u}{\partial t} &= \frac{\partial}{\partial x}\left(\mathcal{D}(u)\frac{\partial u}{\partial x}\right) + \frac{\partial}{\partial y}\left(\mathcal{D}(u)\frac{\partial y}{\partial x}\right) + f(u), && t > t_0,\ \mathbf{x} \in \Omega, \\
\frac{\mathrm{d}u}{\mathrm{d}t} &= f(u), && t > t_0,\ \mathbf{x} \in \partial\Omega, \\
u(\mathbf{x}, t) &= 0 && t = t_0,\ \mathbf{x} \in \Omega, \\
u(\mathbf{x}, t) &= u_0 && t = t_0,\ \mathbf{x} \in \partial\Omega.
\end{aligned}
\tag{S1}
$$

Here, $\Omega$ represents the set of points inside the pore (such that $0 < x < L$ and $0 < y < L$) and $\partial\Omega$ represents the set of points on the pore boundary (such that $x \in \{0, L\}$ or $y \in \{0, L\}$). Due to the symmetry of the problem, we solve the PDE on a quarter of the full domain (Fig. S1*c*) and apply reflecting (symmetric) boundary conditions on the interior boundary.

We used a method of lines based finite difference approach to approximate the solution of the PDE model, $u(\boldsymbol{x}, t)$, with a system of ordinary differential equations (ODEs), $\{u_{i,j}(t)\}_{i,j}$. Our implementation is available in `Module/Functions/SolvePDE.jl` on GitHub. We discretise the domain into a square mesh with $N^2$ nodes, denoting $\mathbf{x}_{i,j} = (x_i, y_j) = ((i-1)\Delta, (j-1)\Delta)$ for $i, j = 1, 2, ..., N+1$ the location of points that surround each element (Fig. S1*d*). This yields

$$
\begin{aligned}
\frac{\mathrm{d}u_{i,j}}{\mathrm{d}t} = &\frac{1}{2\Delta^2}\Big[\big(\mathcal{D}(u_{i,j}) + \mathcal{D}(u_{i+1,j})\big)(u_{i+1,j} - u_{i,j}) - \big(\mathcal{D}(u_{i,j}) + \mathcal{D}(u_{i-1,j})\big)(u_{i,j} - u_{i-1,j})\Big] \\
&+ \frac{1}{2\Delta^2}\Big[\big(\mathcal{D}(u_{i,j}) + \mathcal{D}(u_{i,j+1})\big)(u_{i,j+1} - u_{i,j}) - \big(\mathcal{D}(u_{i,j}) + \mathcal{D}(u_{i,j-1})\big)(u_{i,j} - u_{i,j-1})\Big] \\
&+ f(u_{i,j}),
\end{aligned}
\tag{S2}
$$

which applies on the interior nodes, $i, j = 2, 3, ..., N$ (Fig. S1*d*, grey markers). On the interior boundary where $i = N+1$ or $j = N+1$ (Fig. S1*d*, orange markers) we apply Eq. (S2) and replace $u_{i+1,j} = u_{N-1,j}$ at $i = N+1$, $u_{i,j+1} = u_{i,N-1}$ at $j = N+1$, and $u_{i+1,j+1} = u_{N-1,N-1}$ at $i = N+1$, $j = N+1$. On pore boundary nodes (Fig. S1*d*, blue markers) $i = 1$ or $j = 1$, the boundary condition can be applied exactly

$$
\frac{\mathrm{d}u_{i,j}}{\mathrm{d}t} = f(u_{i,j}). \tag{S3}
$$

We apply the initial condition by setting

$$
u_{i,j}(t_0) = \begin{cases} u_0, & i = 1 \text{ or } j = 1, \\ 0, & \text{otherwise.} \end{cases} \tag{S4}
$$

We integrate the ODE system using the `Tsit5` (a 4th and 5th order Runge-Kutta ODE solver with variable timestep) in `Julia` [1, 2].

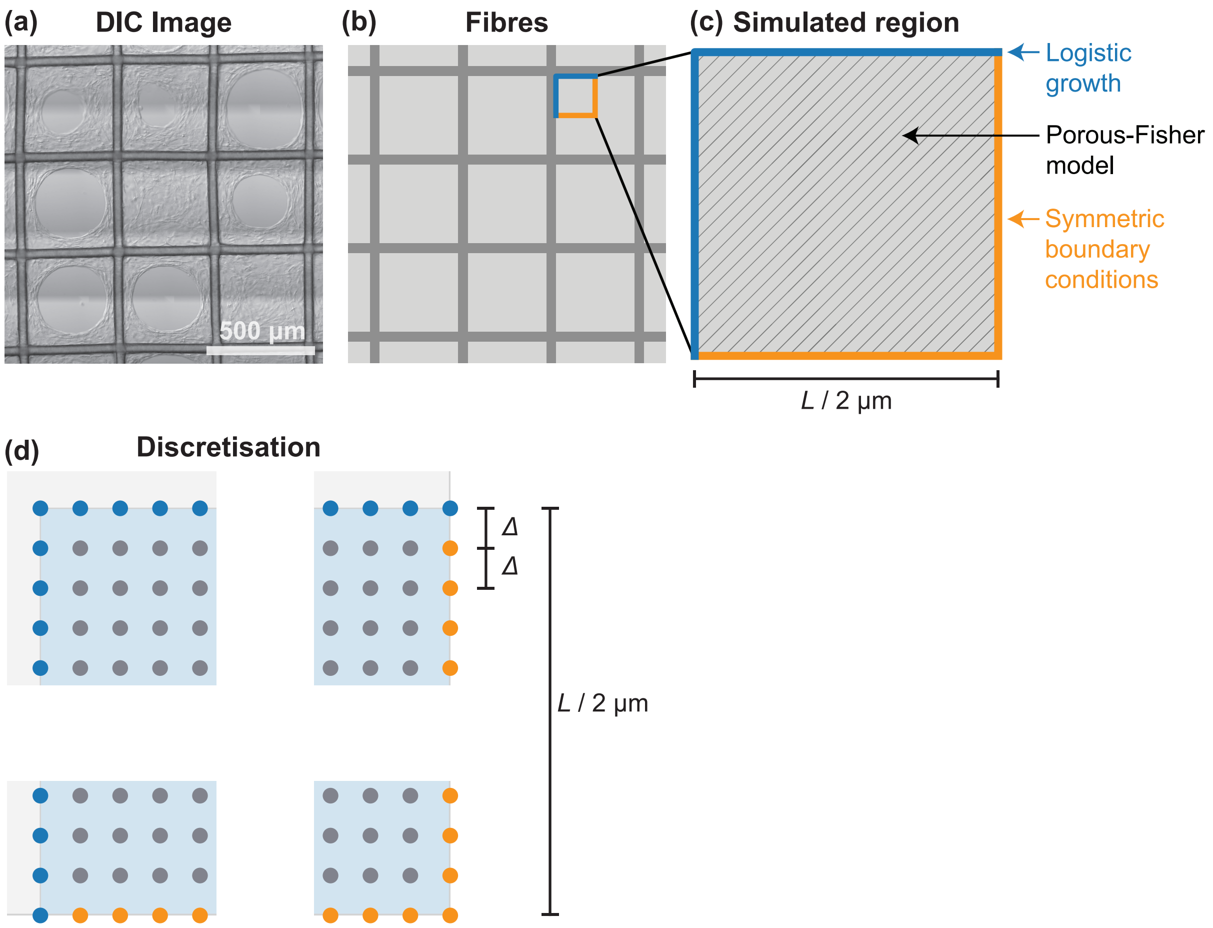


**Figure S1.** **Computational domain and finite difference discretisation.**

#### S2 Interpolating tissue void boundary from PDE solution

It is desirable, to ensure the optimisation routines used to calculate the maximum likelihood estimate converge to a maxima, that all summary statistics are continuous functions of the parameter space, $\boldsymbol{\theta}$. While the numerical solution of the PDE, $\{u_{ij}(t)\}_{i,j=1}^{N}$ varies continuously in $\boldsymbol{\theta}$, the naive approximation of tissue coverage calculated using nodes above the tissue boundary, $\{u_{ij}(t) > \tau K\}_{i,j=1}^{N}$, is not continuous. Therefore, we apply a linear interpolation procedure to estimate the tissue area and perimeter contribution from each finite element surrounded by nodes with tissue densities $\{u_{i,j}, u_{i+1,j}, u_{i,j+1}, u_{i+1,j+1}\}$, which we denote $A(u_1, u_2, u_3, u_4)$ and $P(u_1, u_2, u_3, u_4)$. The area and perimeter of the tissue void, used in calculation of the coverage and circularity summary statistics in the main document, are then given by

$$A_{\text{void}} = 4 \sum_{i=1}^{N-1} \sum_{j=1}^{N-1} A(u_{i,j}, u_{i+1,j}, u_{i,j+1}, u_{i+1,j+1}), \tag{S5}$$

$$P_{\text{void}} = 4 \sum_{i=1}^{N-1} \sum_{j=1}^{N-1} P(u_{i,j}, u_{i+1,j}, u_{i,j+1}, u_{i+1,j+1}). \tag{S6}$$

Note the factor of four as the numerical solution comprises a quarter of each pore.

The interpolation procedure is illustrated in Fig. S2 and is implemented in `Module/Functions/SummaryStatistics.jl`, available on Github. In summary, the number of corners above the threshold, $c_{\text{above}}$, is first determined. The finite element in question is, therefore, represented by Case $(c_{\text{above}} + 1)$ in Fig. S2. In our implementation, we rotate and/or reflect the element to match the configuration of the corresponding case in Fig. S2 exactly. We then linearly interpolate along the edges of the element to approximate the contour $u(t) = 0$. The region containing tissue is then approximated as a polygon (yellow region) formed by joining the interpolated points on the edges with a chord (black-dashed line). The area of this polygon, and the length of the chord, approximate the area and perimeter contribution from this element, giving $A(\cdot)$ and $P(\cdot)$.

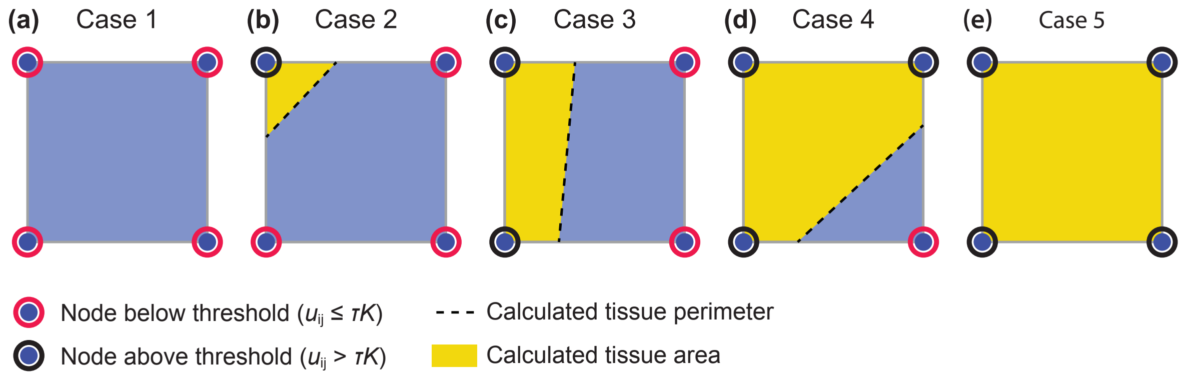


**Figure S2. Interpolation procedure to calculate the area covered by tissue and the perimeter of the tissue void.**

#### S3 Noise model

We pre-estimate a noise model [3] of the form

$$\sigma_i(\mu_i) = \max(0, a_i\mu_i^2 + b_i\mu_i + c_i), \tag{S7}$$

using the observed group standard deviations in the experimental data, pooled for all pore sizes Fig. S3. In Eq. (S7), $\sigma_i(\mu_i)$ is the standard deviation for the $i$th summary statistic, where $\mu_i$ is the mean (equivalent to the model prediction). In all cases, we fix the intercept $c_i$ equal to 10% of the maximum standard deviation observed in the experimental data, and estimate $a_i$ and $b_i$ by minimising the sum of squares of the residual.

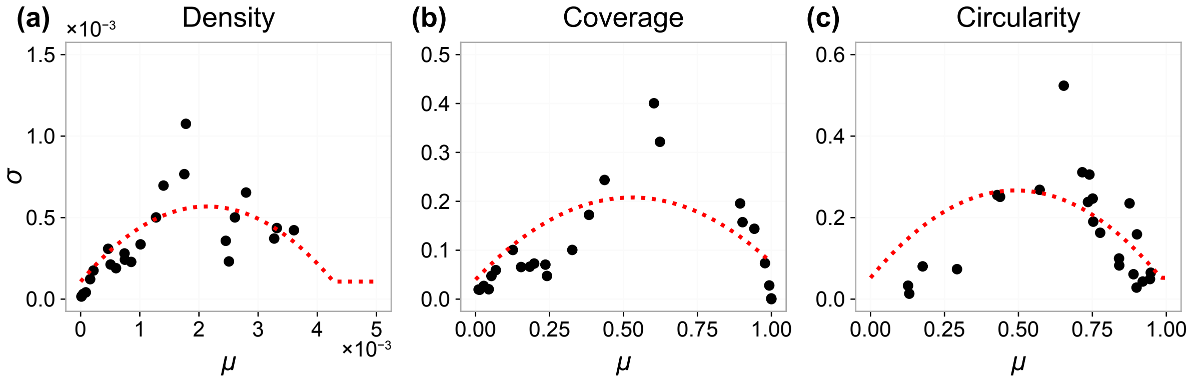


**Figure S3. Pre-estmated noise model.** Observed group standard deviations (black markers) and fitted noise model (red dashed).

#### S4 Including all $L = 400\,\mu\mathrm{m}$ data

In Fig. S4 we repeat results from the main paper for the $400\,\mu\mathrm{m}$ pores in the case that all $400\,\mu\mathrm{m}$ data is included. We see results inconsistent with the inferred results for the other pore sizes, seeing very low estimates for the proliferation rate, $\lambda$. When we exclude data from $t = 7\,\mathrm{d}$ for the main results, we see estimates for the proliferation rate that are similar to the other pore sizes, suggesting that these experimental replicates may be outliers. As scaffolds are fixed prior to staining and imaging, data from these scaffolds at earlier times is not available, and it is, therefore, difficult to ascertain why these scaffolds may exhibit behaviour different to the others.

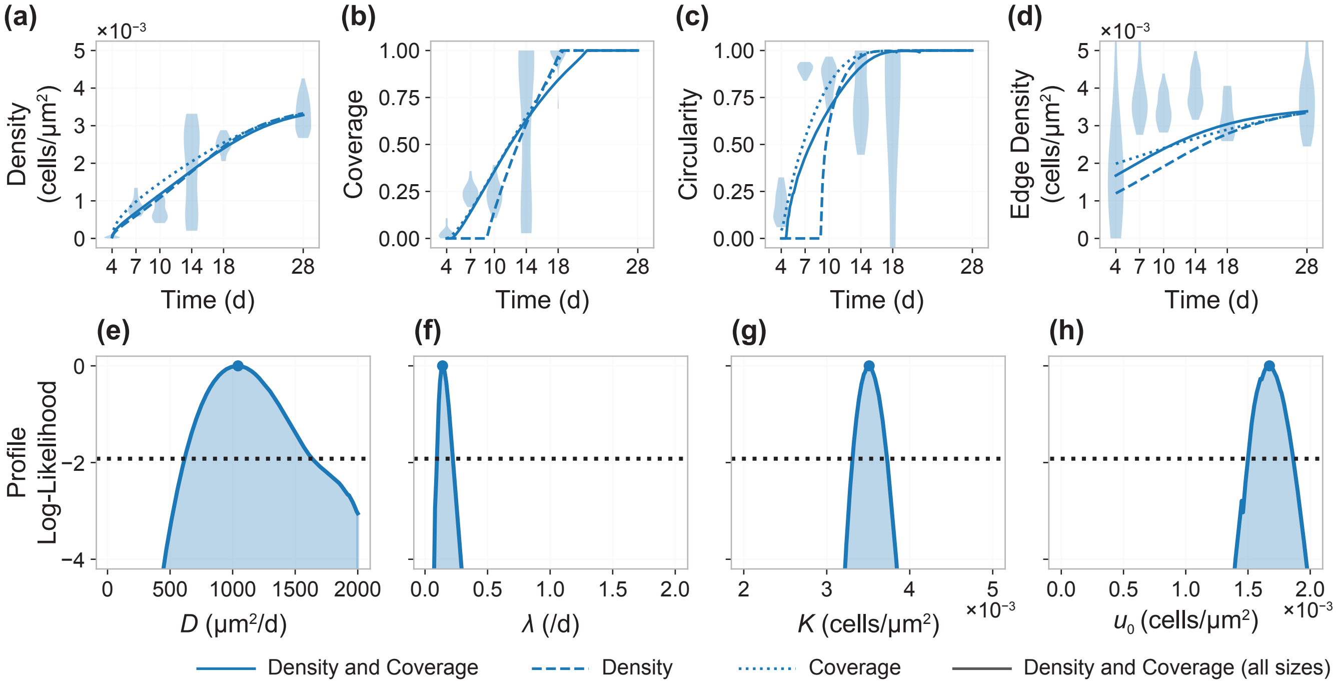


**Figure S4. Results including all $L = 400\,\mu\mathrm{m}$ data.** (a–d) Violin plots show the experimental data. In each case, a model prediction is shown based on the maximum likelihood estimate where the cell density (dashed colour); cell density and tissue coverage (solid colour); tissue coverage with day 28 density measurement (dotted colour); and cell density and tissue coverage from all pore sizes (solid grey). (e–h) Profile likelihoods for each inferred parameter where both the density and coverage are used. Dotted horizontal black line indicates the $-1.92$ contour which corresponds to an asymptotic 95% confidence interval for each parameter.

#### S5 Inference using circularity

In Fig. S5 and Fig. S6 we repeat results from the main paper where the circularity is included alongside cell density and tissue coverage information. We see that, even when we calibrate the model to circularity data, we still do not match the circularity measurements seen in the experimental data, in particular for the case where inference is performed on data from all pore sizes simultaneously.

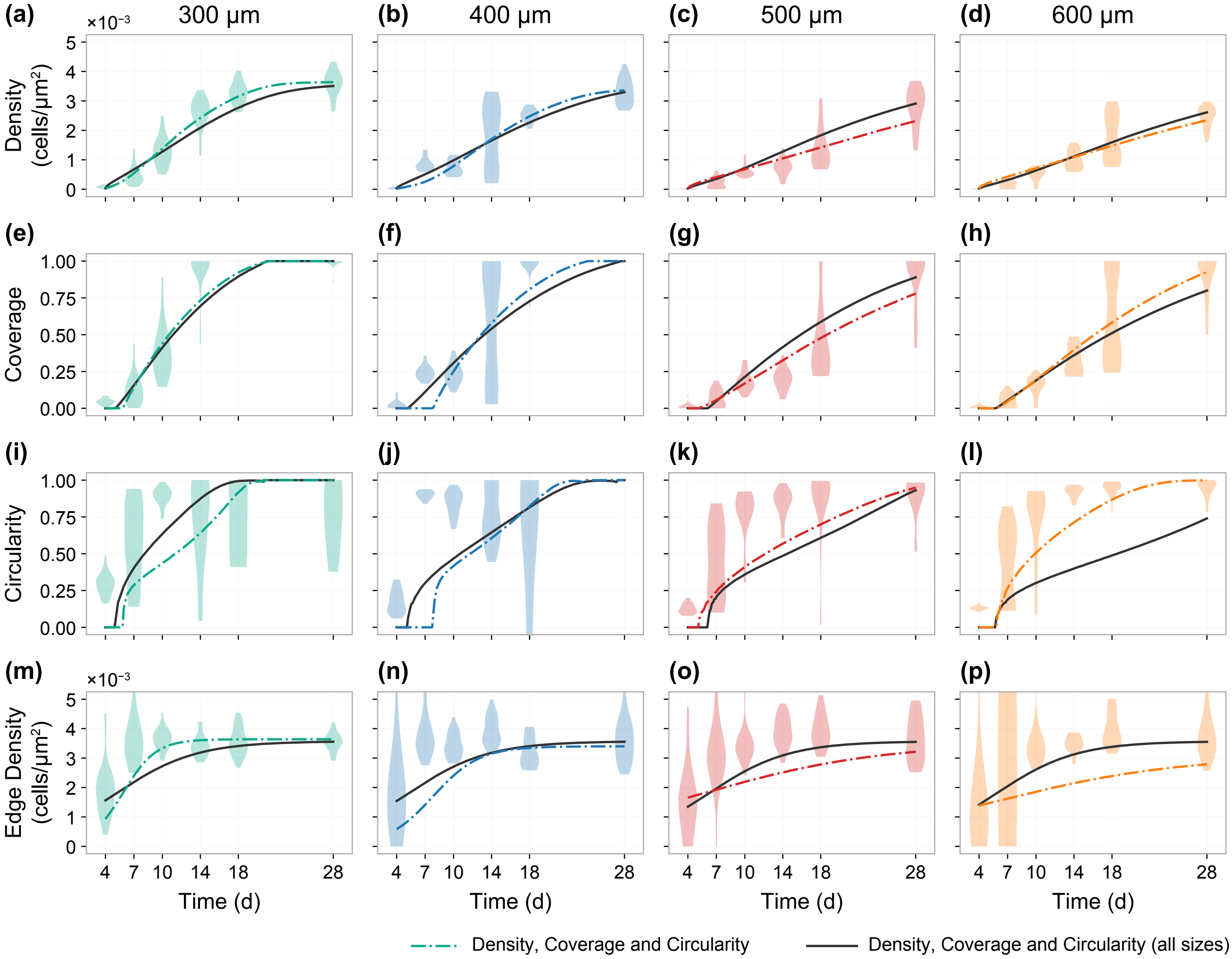


**Figure S5. Processed experimental data and model fits where the circularity is also used for inference.** Experimental data and model fit showing (a–d) the density (cells/μm$^2$), (e–h) the coverage, (i–l) the circularity, and (m–p) the edge density. Violin plots show the experimental data. In each case, a model prediction is shown based on a maximum likelihood estimate that includes the cell density, tissue coverage, and tissue void circularity from (coloured) each individual pore size; and, (grey) simultaneously from all pore sizes.

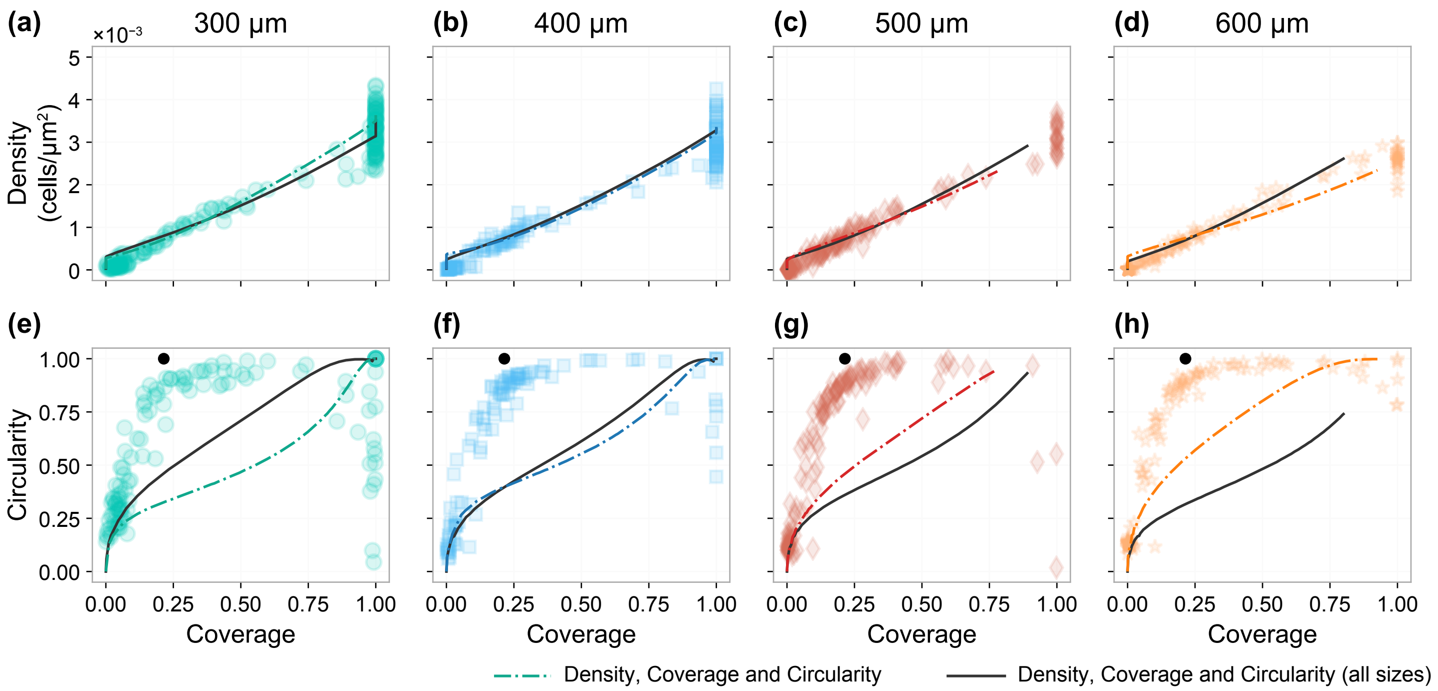


**Figure S6. Correlations between experimental and simulated summary statistics where the circularity is also used for inference.** Experimental data and model fit showing the relationship between (a–d) tissue coverage and cell density, and (e–h) coverage and circularity. In each case, a model prediction is shown based on a maximum likelihood estimate that includes the cell density, tissue coverage, and tissue void circularity from (coloured) each individual pore size; and, (grey) simultaneously from all pore sizes.

#### S6 Fisher-Kolmogorov model

In the main text, we note that cells cannot move freely on the experimental substrate. Rather, cells work together to bridge the pore through interconnecting material such as extracellular matrix and intracellular actin filaments. This assumption leads to a governing equation of the form

$$\frac{\partial u}{\partial t} = D\nabla \cdot \left[ \left( \frac{u}{K} \right)^{\alpha} \nabla u \right] + \lambda u \left( 1 - \frac{u}{K} \right), \quad \mathbf{x} \in \Omega, \tag{S8}$$

where $\alpha = 1$, often referred to as the *Porous-Fisher equation.* Here, we relax the assumption of collaborative cell migration and reproduce the main results in the case where $\alpha = 0$, i.e., cell migration is a linear diffusion process. This choice recovers what is commonly referred to as the *Fisher-Kolmogorov model* [4], however does not align with the collaborative migration process that gives rise to tissue growth in the experiments, and will not yield a sharp-fronted tissue interface.

In Fig. S7 and Fig. S8, we compare experimental data with model best fits from the Fisher-Kolmogorov model. In Fig. S9 we show profile likelihoods for this model. As with results in the main paper, we find we are unable to match features in the experimental data relating to circularity. However, as we are able to account for both cell density and tissue coverage, we are able to quantify cell proliferation rates for each parameter, which we find to be similar to results in the main document.

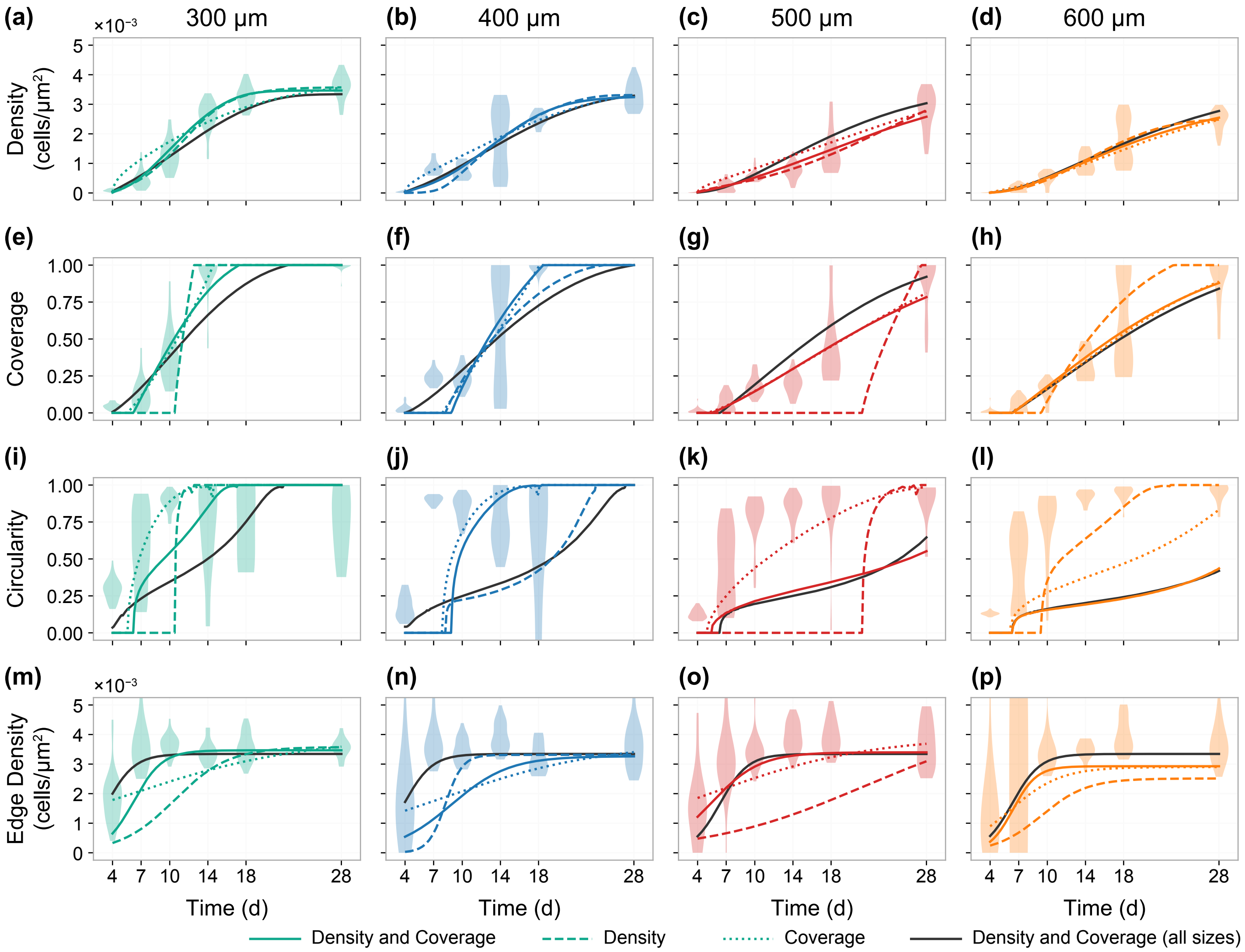


**Figure S7. Processed experimental data and model fits using the Fisher-Kolmogorov model.** Experimental data and model fit showing (a–d) the density (cells/μm$^2$), (e–h) the coverage, (i–l) the circularity, and (m–p) the edge density. Violin plots show the experimental data. In each case, a model prediction is shown based on the maximum likelihood estimate where the cell density (dashed colour); cell density and tissue coverage (solid colour); tissue coverage with day 28 density measurement (dotted colour); and cell density and tissue coverage from all pore sizes (solid grey).

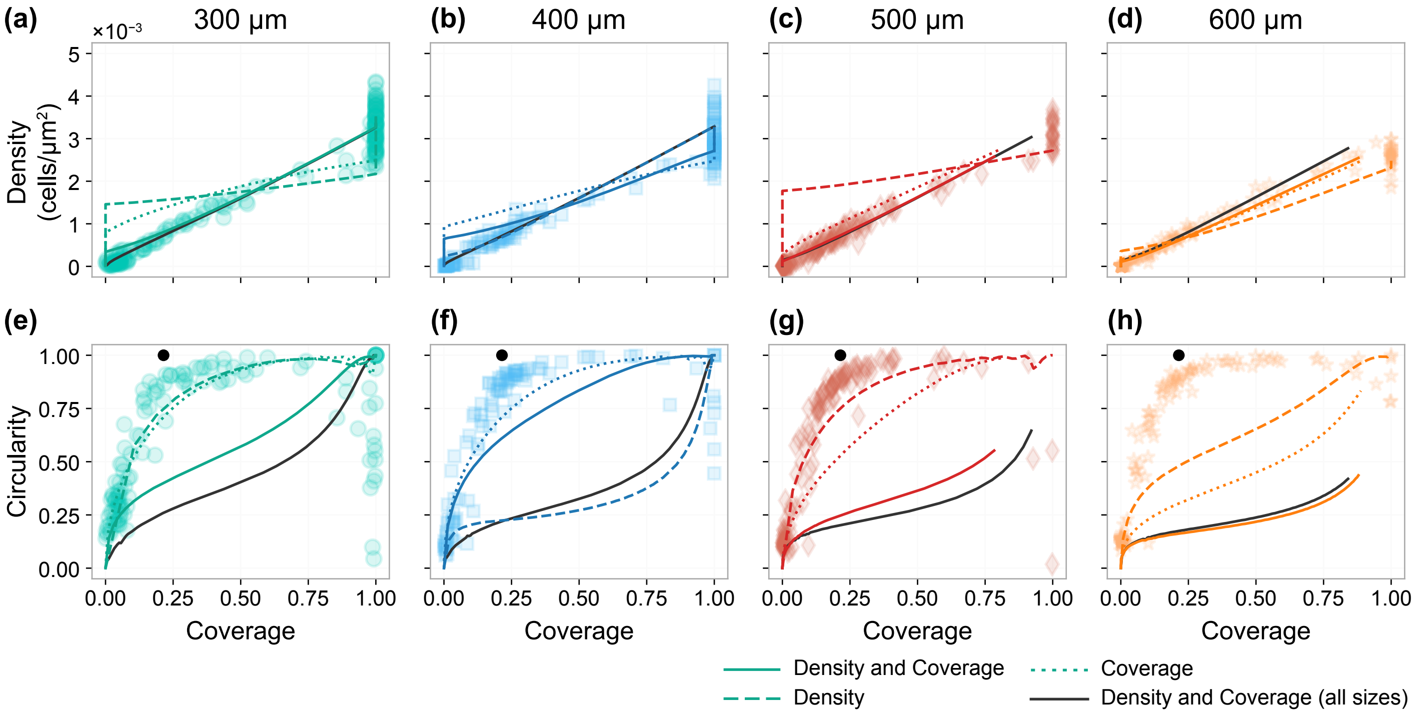


**Figure S8. Correlations between experimental and simulated summary statistics using the Fisher-Kolmogorov model.** Experimental data and model fit showing the relationship between (a–d) tissue coverage and cell density, and (e–h) coverage and circularity. In each case, a model prediction is shown based on the maximum likelihood estimate where the cell density (dashed colour); cell density and tissue coverage (solid colour); tissue coverage with day 28 density measurement (dotted colour); and cell density and tissue coverage from all pore sizes (solid grey).

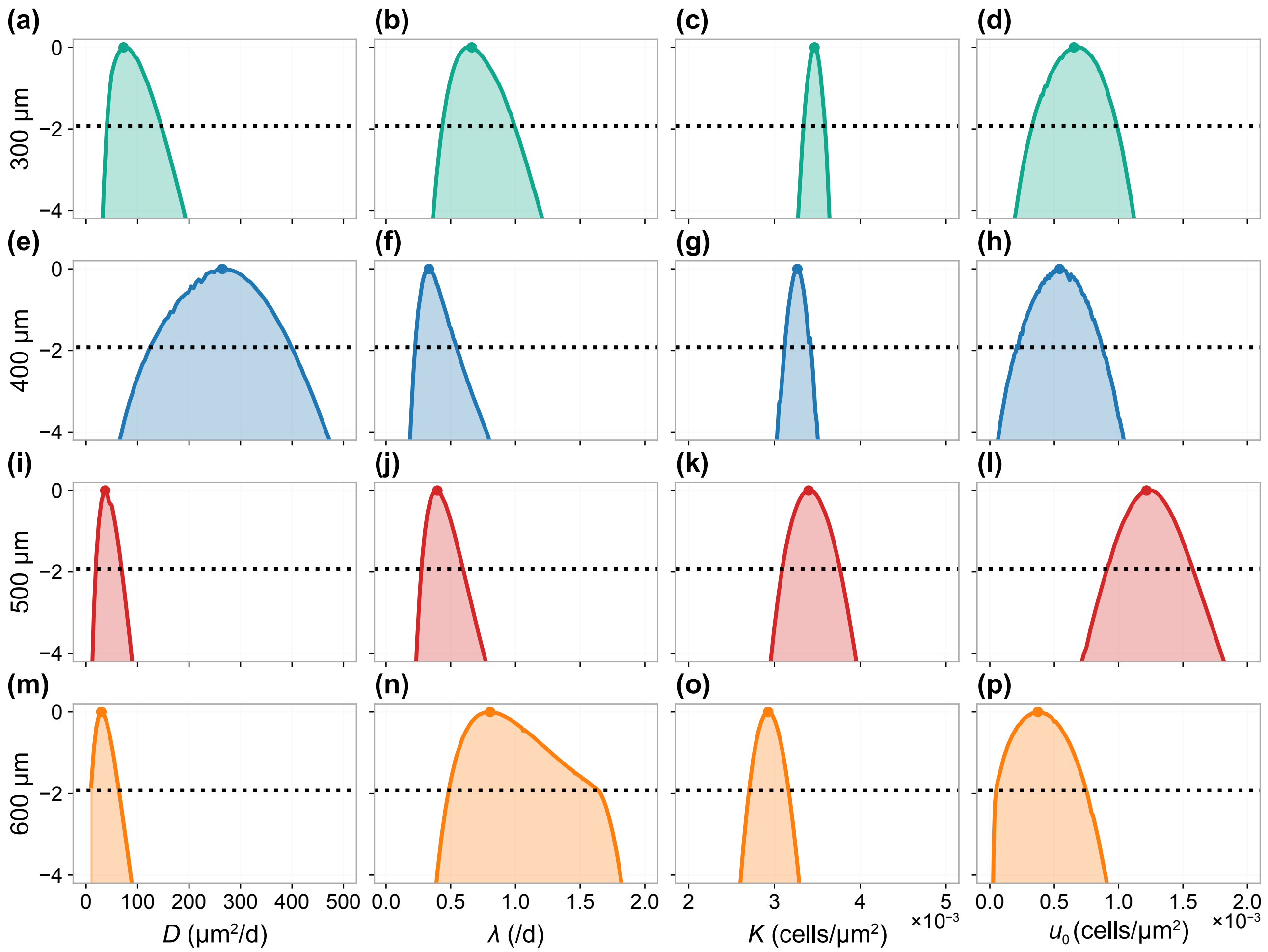


**Figure S9. Correlations between experimental and simulated summary statistics using the Fisher-Kolmogorov model. Profile likelihoods for calibrated model parameters.** Profile likelihoods for each inferred parameter and pore size where only the density is used (dashed) and where both the density and coverage are used (solid). Dotted horizontal black line indicates the $-1.92$ contour which corresponds to an asymptotic 95% confidence interval for each parameter. Also shown are profiles for $D$, $\lambda$ and $K$ where information relating to cell density and tissue coverage information from all pores is included simultaneously.
